# A Syntenic Pangenome of *Gardnerella* Reveals Novel Plasmids and Phage, Taxonomic Boundaries, and Species-Level Stratification of Metabolic and Virulence Potential

**DOI:** 10.1101/2025.02.19.636902

**Authors:** Heather K. Bouzek, Martha A. Zepeda-Rivera, Sujatha Srinivasan, Elliot M. Lee, Susan M. Strenk, Dakota S. Jones, Elsa F. McMahon, Tina L. Fiedler, Marko Kostovski, Michael T. France, Jacques Ravel, David N. Fredricks, Christopher D. Johnston

## Abstract

*Gardnerella* species are central to bacterial vaginosis (BV), a condition affecting nearly one in three women of reproductive age and associated with preterm birth and increased susceptibility to sexually transmitted infections. Despite decades of study, progress in defining *Gardnerella* diversity has been hindered by inconsistent taxonomy and poor-quality genomic resources. Here, we sequenced 392 new *Gardnerella* isolates from asymptomatic and BV-associated microbiota and integrated this collection with all publicly available genomes. After stringent curation, we generated a high-quality reference set of 313 distinct genomes that underpins a comprehensive taxonomic framework. Using average nucleotide identity (ANI), digital DNA–DNA hybridization (dDDH), and phylogenomics, we resolved 21 genomic lineages encompassing 11 species and 15 subspecies, each assigned a provisional formal name. Integration of complete long-read assemblies enabled construction of the first syntenic *Gardnerella* pangenome, revealing lineage-specific repertoires of virulence, metabolic, and defense systems - including variable sialidases (NanH), vaginolysin, and amino-acid biosynthetic pathways - and defining conserved genomic architecture across species. Comparative methylome profiling further highlighted restriction-modification system diversity that may influence genetic exchange. Finally, we identified the first native cryptic plasmids in *Gardnerella*, overturning the assumption that the genus lacks plasmids, and demonstrated their use in generating a replicative *E. coli*-*Gardnerella* shuttle vector. Together, these results establish a complete genomic and functional framework for *Gardnerella*, providing a reproducible foundation for mechanistic and translational studies of BV and a model for resolving taxonomy and functional stratification in other urogenital-associated bacteria.

## Main

Bacterial vaginosis (BV) is a common and global condition characterized by an imbalance in the vaginal microbiota^1–3^, associated with significant health risks, including increased susceptibility to sexually transmitted infections^4,5^ and adverse pregnancy outcomes such as preterm birth and low birth weight^6–8^. While molecular methods have associated many bacterial species with BV, *Gardnerella vaginalis* was the first bacterium isolated from individuals affected with BV^9^, and was long thought to be the sole causative agent of this condition^1,10,11^ due to its high virulence potential^3,12–20^, its presence in 95–100% of BV cases^11,21,22^ and its ability to thrive in the vaginal environment^10,19,20,23–25^. Our previous research revealed that while *Gardnerella vaginalis* is correlated with a greater risk ratio for BV than other validly named *Gardnerella* species, no single *Gardnerella* species is specific for BV based on a *cpn60* qPCR. In fact, the presence of a high *Gardnerella* species diversity was identified as a risk factor for the development of BV in our longitudinal study^22^. The presence of *Gardnerella* species in both BV-affected and asymptomatic individuals suggests the potential for both commensal and pathogenic lineages or states^3^. However, the genetic capabilities of individual *Gardnerella* lineages remain unclear, partially due to inconsistent taxonomic assignments and overly complex *Gardnerella* nomenclature^19^, despite laudable efforts from multiple research groups^26–35^. Further, many of the public *Gardnerella* genome assemblies are duplicated, contaminated, or incompletely assembled and did not pass our quality metrics for pangenome analysis. Thus, we sequenced over 392 *Gardnerella* isolates genomes from both asymptomatic individuals or those with BV and combined these with all available public genomes to generate a highly curated and deduplicated collection for analyses. We applied comprehensive comparative genomic analysis of this curated collection to taxonomically identify *Gardnerella* lineages and propose a single naming scheme that is reflective of their phylogenetic placement and historical context. Syntenic pangenome partitioning revealed lineage-specific repertoires of metabolic pathways, virulence-associated genes, and defense systems. Collectively, this study integrates pangenome analyses, genome-based taxonomy, and experimental validation to define species boundaries within *Gardnerella*, characterize lineage-specific metabolic and virulence-associated potential, and identify mobile genetic elements, including a cryptic plasmid that enables the development of a shuttle vector for the genetic engineering of *Gardnerella*.

## Results

### Gardnerella and Bifidobacterium are distinct genera

We obtained genomes for 392 *Gardnerella* isolates (Fredricks and Ravel bacterial isolate collections) from asymptomatic individuals and those with BV. We curated these and additionally available public *Gardnerella* genomes (n=271 within the NCBI database as of May 2025) using stringent quality metrics to remove duplicate and low-quality genomes (Methods, Supplementary Table 1). This resulted in a final curated set of 312 genomes for comparative genomic analyses (Supplementary Table 2). Analysis of the *cpn60* gene (which encodes a 60 kDa chaperonin) currently provides the most reliable method for *Gardnerella* species typing, as these species are indiscernible by traditional 16S rRNA gene sequence typing^36,37^. BLASTn analysis of the *cpn60* gene of these isolates supported their identification as *Gardnerella* species (Supplementary Table 2).

Despite the classification of these genomes as *Gardnerella* against and within the NCBI database, initial speciation against the Genome Taxonomy Database (GTDB) assigned many of these genomes to *Bifidobacterium* (Supplementary Table 2). However, *Bifidobacterium* is known to be a polyphyletic group, consisting of at least six distinct clades^38,39^, whose members exhibit substantial genetic diversity. The separation of *Gardnerella* from *Bifidobacterium* has previously been demonstrated through comparisons of genome length, G+C percentage, and overall genome composition^31,39^. Consistent with theses prior observations, comparisons of genome length and G+C percentage between our *Gardnerella* genome set, *Bifidobacterium,* and closely related scardovial genera (e.g., *Alloscardovia*^40^*, Neoscardovia*^41^*, Aeriscardovia*^42^, *Scardovia*^43^) (n=229 genomes; Supplementary Table 3) indicate that *Gardnerella* genomes are significantly shorter (p<0.001, Wilcoxon *post hoc* test) and have lower G+C content (p<0.001, Wilcoxon *post hoc* test) than other *Bifidobacterium* members (Extended Data Fig. 1a-b). Notably, *Gardnerella* shares a human-focused and pathogenic niche with *Alloscardovia, Aeriscardovia* and *Scardovia* and clusters more closely with the scardovial genera across these genomic features than with *Bifidobacterium.* Additionally, average amino acid identity (AAI) values between *Gardnerella* and *Bifidobacterium* fell below genus thresholds of 65%^44^ (Extended Data Fig. 1c). Together, these results confirmed that our final curated genome set comprises only *Gardnerella* species, further supporting previous evidence that *Gardnerella* and *Bifidobacterium* represent distinct genera within the *Bifidobacteriaceae* family.

### Identification and proposed revised nomenclature for *Gardnerella* lineages

Given the importance of *Gardnerella* species diversity in BV^21^, we first assessed the number of lineages present in our genome collection. *Gardnerella* has been classified using various approaches^30–33,45,46^, but consistent naming has been a challenge for *Gardnerella* despite extensive efforts^26–28,32^ (Fig. 1). Early subtyping by Piot *et al*.^26^ relied on sugar hydrolysis phenotypes and later expanded to 17 biotypes^27^. Genomic methods such as restriction fragment length polymorphism (RFLP)^28^ and amplified ribosomal DNA restriction analysis (ARDRA)^29^ identified several distinct genotypes. PCR and whole-genome sequence comparisons introduced two new naming schemes including designations “A - D”^37^ or “1 - 4”^30^ which led to hybrid labels (“C/1”, “B/2”, “A/4”, and “D/3”)^47^. These species were then reframed as reassorted ecotypes^31^ incorporating the genomic context of the distinct types. With widespread whole-genome sequencing, additional standards based on ANI emerged, describing thirteen genomic species (Gsp)^32^, nine genomospecies (GS)^33^, or nine different genotypes (GGtypes)^35^. Subsequent formal naming of a subset of these species included *G. vaginalis*^9^, *G. pickettii*^48^, *G. piotii*^32^, *G. leopoldii*^32^, *G. swidsinskii*^32^, *G. greenwoodii*^48^, *G. lacydonensis*^49^, *G. bretellae*^49^, *G. massiliensis*^49^, and *G. phocaeensis*^49^ (Fig. 1). Despite the use of diverse classification schemes, phylogenetic analyses across studies consistently recover a deep bifurcation within *Gardnerella*, with the same two major lineages forming the basal split of the genus. While this division has been explicitly designated as “Set A” and “Set B” only by Ahmed *et al*., analogous groupings confirm the primary branching structure in multiple independent phylogenetic frameworks^30–35,37,46,50,51^.

**Figure 1:**
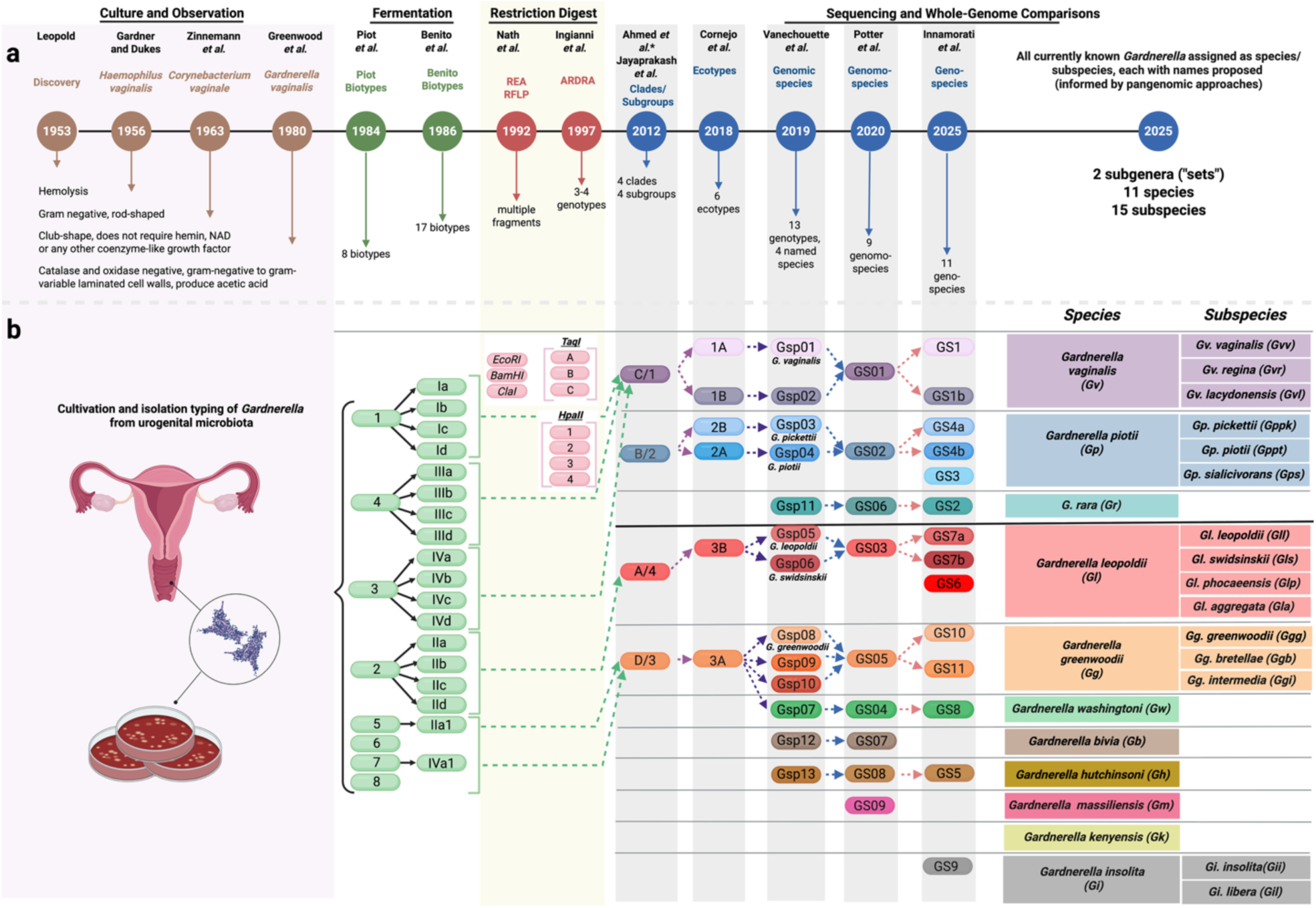
Historical context of *Gardnerella* nomenclature. **a**, From top to bottom, the schematic shows the identification method by type, publication first author, names given to *Gardnerella* or naming framework, date of publication, and resulting conclusions from naming schemes. **b**, *Gardnerella* subtyping resulting from the publication in (**a**), where each column represents a subset of *Gardnerella*. Arrows indicate consistent lineages between studies without implying a definitive link. The two right-most columns contain updated lineage assignments with corresponding species and subspecies nomenclature proposed within this study.

Consistent with this, *cpn60* gene sequences and reference-free whole-genome phylogenies separated all genomes in our collection into two deeply divergent lineages (Fig. 2, Extended Data Fig. 2, Supplementary Table 4) here maintaining the Set A and Set B designation. To determine the number of *Gardnerella* species represented, we calculated pairwise genome-wide average nucleotide identity (ANI) values across all genomes and examined their distribution (Extended Data Fig. 3a). In accordance with prior large-scale analyses demonstrating a pronounced discontinuity in ANI values around the canonical prokaryotic boundary, most inter-lineage comparisons fell well below 95%, whereas closely related genomes clustered above 95% with an exceedingly small fraction of the sequence comparisons falling above 94% (Extended Data Fig. 3a). Because ANI and digital DNA–DNA hybridization (dDDH) values were strongly colinear across the dataset, dDDH did not provide additional resolution but independently supported the same groupings: all genomes that clustered together by ANI also exceeded the 70% dDDH species threshold (Extended Data Fig. 3b–c). Thus, species boundaries inferred from ANI were fully congruent with the universally accepted 70% dDDH. Integrating whole-genome phylogenies with ANI/dDDH clustering, we identified eleven *Gardnerella* species. Within five of these species, additional phylogenomic structure supported sub-species level differentiation, resulting in twenty-one distinct *Gardnerella* lineages (Fig. 2, Supplementary Table 4). These species-level assignments were congruent with the classification scheme proposed by Potter^33^. The *Gardnerella* species *bretellae, lacydonensis, leopoldii, phocaeensis*, *pickettii, piotii*, and *swidsinskii* were previously described as distinct species when integrating phenotypic characterization with genome-based analyses. In our dataset, these lineages likewise form well-supported monophyletic clades in whole-genome phylogenies and remain clearly differentiated from other *Gardnerella* taxa. However, when evaluated across the expanded genome collection as analyzed here, pairwise ANI and dDDH comparisons reveal that these lineages consistently fall within the genomic coherence of broader species-level clusters defined by ≥95% ANI and ≥70% dDDH thresholds. While they are phylogenetically structured and phenotypically distinguishable, they do not cross the currently accepted genomic discontinuity thresholds that would necessitate recognition as separate species under genome-based standards.

**Figure 2:**
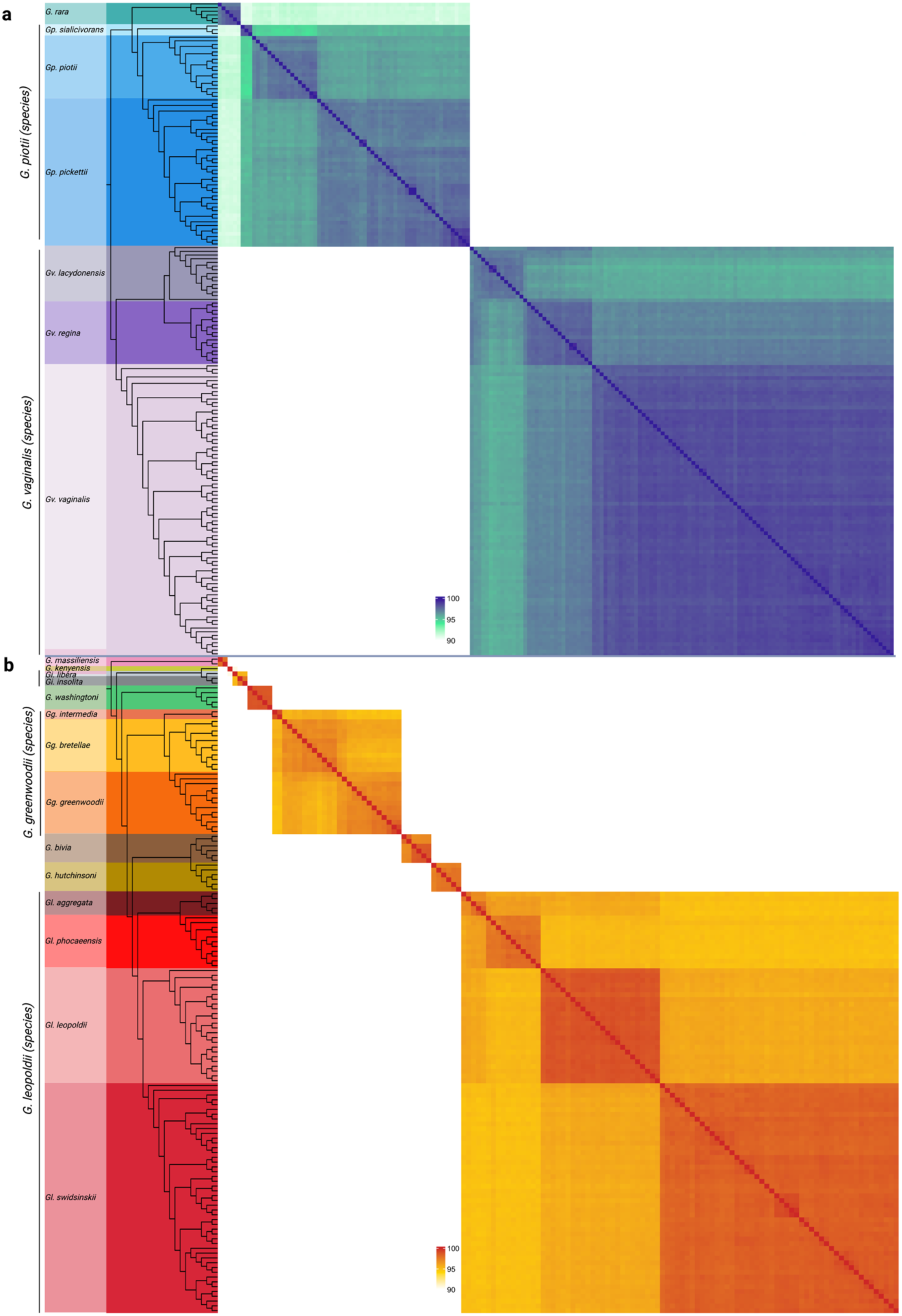
Average nucleotide identity values between *Gardnerella* genomes. Average nucleotide identity values between *Gardnerella* genomes in **a**, Set A and **b**, Set B. Key for each panel indicates corresponding values to each color. White represents values below 90%. Genomes are organized based on a kSNP4-generated whole-genome reference-free phylogeny. Figure is a composite of the kSNP4 tree created in iTOL and the map sorted by the values in the kSNP4 tree as created using ComplexHeatmap in R.

Given the extensive history of overlapping and partially incompatible classification schemes for *Gardnerella* (Fig. 1), we next harmonized existing species and subspecies names and established a new nomenclature across all identified lineages (Fig. 1, Fig. 2, Supplementary Tables 2 and 4). Details on the rationale for each proposed nomenclatural revision, including species-specific functional attributes and ecological isolation sources, are provided in the Protologues. Our aim was to integrate previously used species and subspecies names to maintain a consistent framework. The historically recognized “*G. vaginalis*” corresponds to a subspecies-level clade within a broader species and is designated here as *G. vaginalis* subspecies *vaginalis*, preserving historical continuity while reflecting its phylogenetic placement (Fig. 1, Supplementary Table 1, Supplementary Table 4). Similarly, we propose distinct species names for lineages with well-supported genomic and phenotypic divergence, including *G. bivia*, *G. hutchinsoni*, and *G. washingtoni*, among others (Extended Fig. 3g). To minimize ambiguity in referring to *Gardnerella* species in this paper and future efforts, we intentionally propose species names that can be abbreviated with unique two letter abbreviations (*Gb*, *Gg, Gh, Gi*, *Gk*, *Gl*, *Gm, Gp, Gr*, *Gv* and *Gw*), while individual subspecies were designated with discernable three-letter abbreviations (except for *Gardnerella piotii* subsp. *pickettii* and *Gardnerella piotii* subsp. *piotii,* which each require four letters, *Gppk* and *Gppt*) (Fig. 1, Supplementary Table 4).

### A syntenic pangenome for *Gardnerella*

Having stringently identified *Gardnerella* species and subspecies lineages in our genome set and assigning names that adhered as closely to precedent, where possible, we were next able to investigate the genetic features associated with each lineage. Pangenome analysis provides a comprehensive view of the genetic repertoire within a group of bacteria by identifying genetic features that are well-conserved across most (>95%) genomes (“persistent” or “core” genome) versus those that are only present in a subset of genomes (“cloud” or “accessory” genome). While the core genome often contains genes involved in essential cellular processes^52,53^, analysis of the accessory genomes can provide key insights about genetic diversity, adaptation, and evolutionary relationships. For example, previous pangenome efforts in *Gardnerella* have expanded taxonomic groupings, identified metabolically essential genes, and uncovered virulence and niche adaptation features^34,50,54^. However, bacterial genes often function in co-regulated units or clusters involved in specific biological processes (e.g., operons). To identify putative operons within the *Gardnerella* pangenome, we applied a syntenic approach that accounts for gene order by using the Partitioned Pangenome Graph of Linked Neighbors (PPanGGOLiN) tool^52^ (Figs. 3-4, Supplementary Table 5). Briefly, in these analyses, the coding sequences of genes are grouped into “nodes” based on amino acid similarity and visualized as circular nodes, with node size reflecting the relative presence of that gene group across the genome set (Fig. 4a-c). Nodes representing neighboring genes are linked into syntenic units called “modules” (Extended Data Fig. 4). This analysis clearly distinguished between the core (“persistent” partition) and accessory genomes, and enabled identification of accessory genetic elements present at intermediate frequency (“shell” partitions) or low frequency (“cloud” partition) (Fig. 3). By leveraging modules and pangenome partitions, conserved arrangements of genes can be traced across multiple genomes and enables insight into the functional and evolutionary dynamics driving diversity at the genus, species, and strain levels.

**Figure 3:**
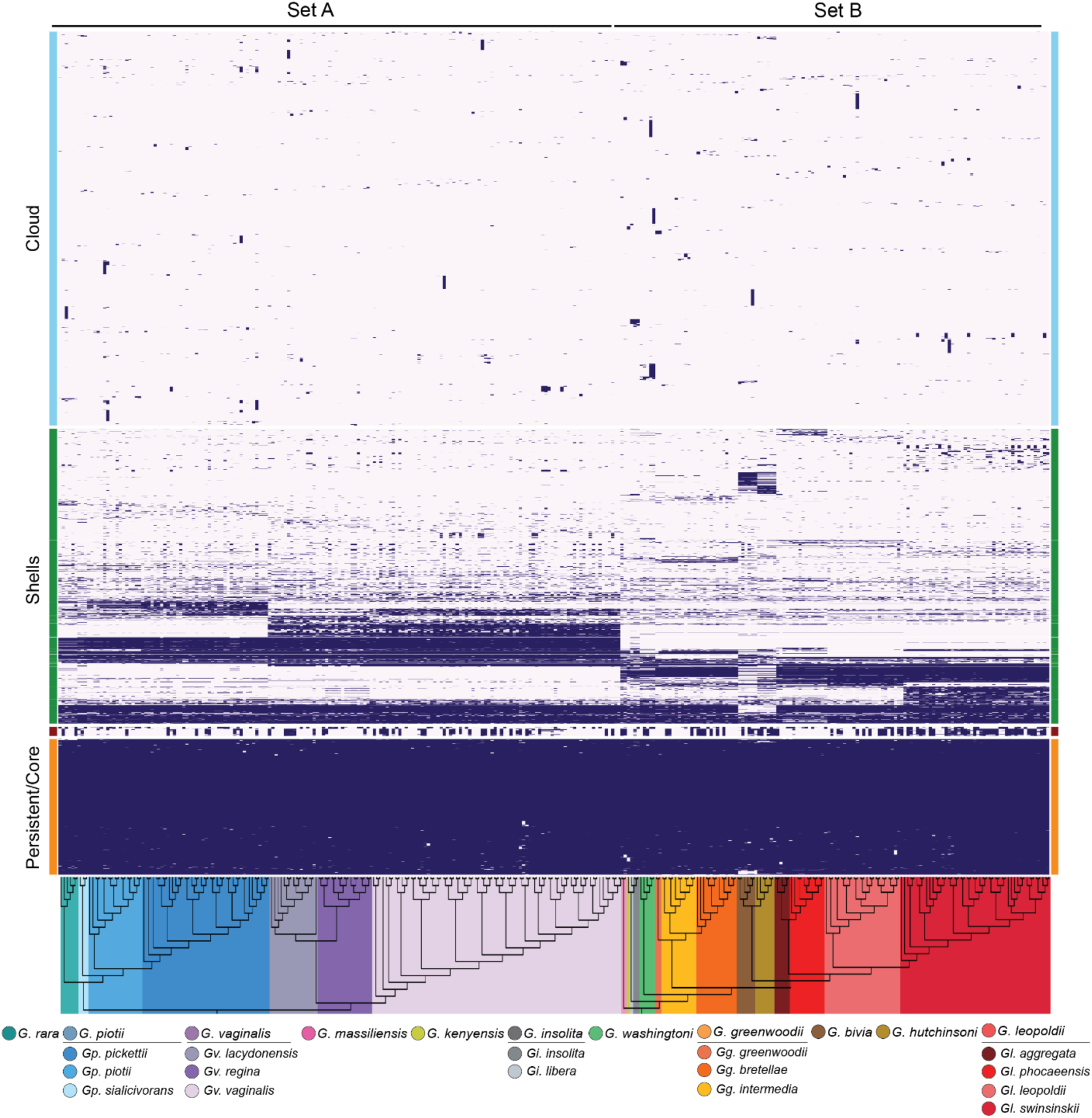
*Gardnerella* pangenome partitions. The tile plot displays the presence and absence of nodes in each partition where each row represents a node, and each column corresponds to a *Gardnerella* genome. Colors on the side correspond to core/persistent (orange), phage (red), shells (green), and cloud (blue) partitions. Genomes are ordered by kSNP4 phylogenies of Set A and Set B, with colors on dendrogram indicating lineage.

**Figure 4:**
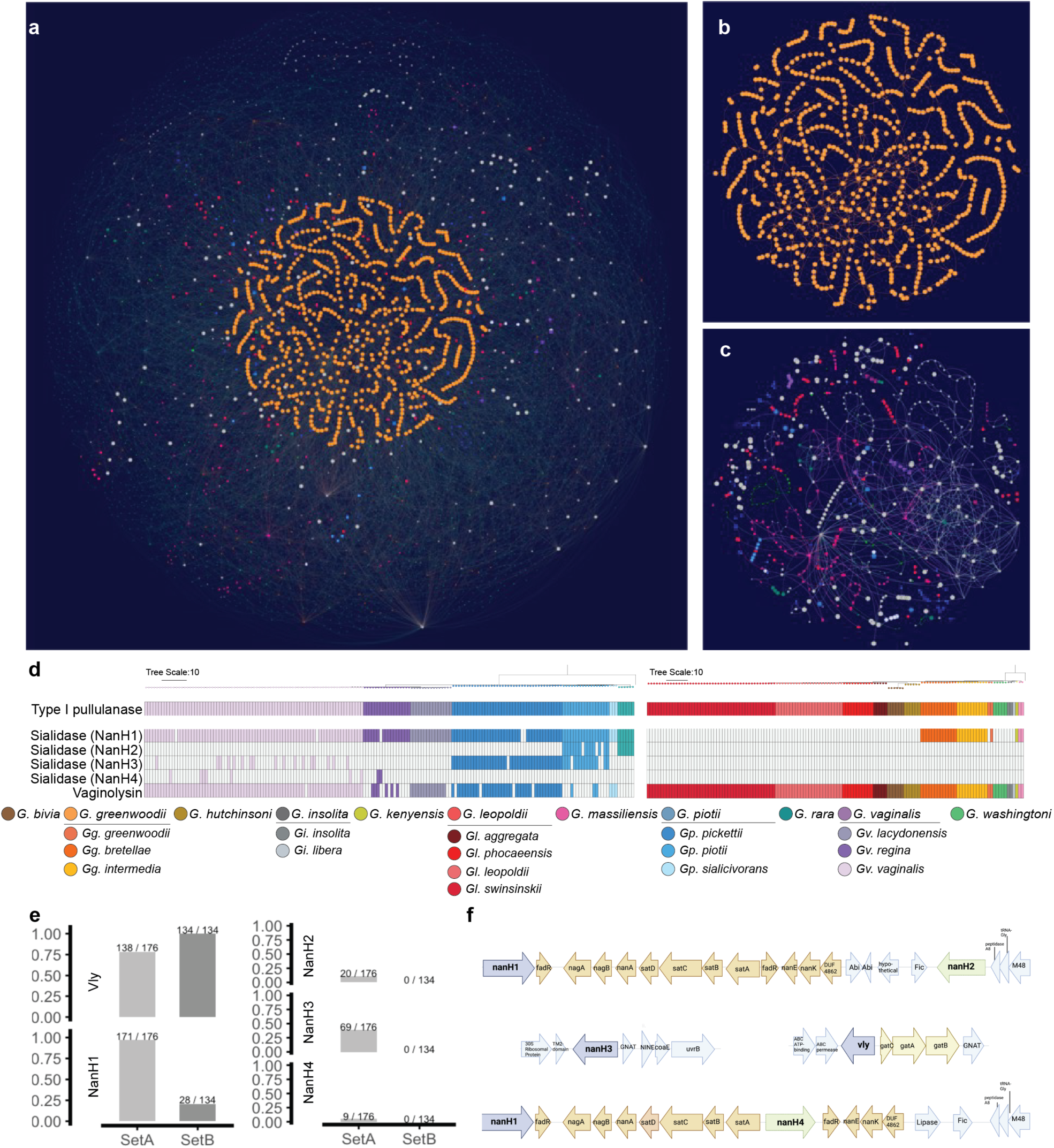
Syntenic maps of the *Gardnerella* pangenome. **a–c,** Visualizations of the PPanGGOLiN-generated *Gardnerella* pangenome. Each node represents a gene group, with edges connecting syntenic neighboring genes. Node size reflects the relative frequency of the gene group across *Gardnerella* genomes. Colors indicate core (orange) or shell (other colors) partitions. **b** and **c**, The core and shell partitions are shown individually. **d**, map of the presence and absence of the sialidase genes (*nanH1*-*nanH4*), type II α-amylase-pullulanase and vaginolysin as colored by the species and subspecies of *Gardnerella*. **e**, Presence–absence bar chart showing the percentage of conserved *Gardnerella* vaginolysin and the sialidase genes across each *Gardnerella* set (gray). Raw presence/absence counts are shown above each bar. **f**, Gene context maps for the sialidase loci.

In our collection, the core genome of *Gardnerella* reflected approximately 16% of identified nodes, enriched in essential cellular functions including translation and ribosomal structure, transcription, RNA processing, signal transduction, energy production, and metabolism of nucleotides, amino acids, inorganic ions, and lipids (Fig. 3, Supplementary Table 5). Periodically, these conserved nodes are interrupted by shell-associated nodes (Fig. 4a; nodes in distinct colors, branching from orange nodes), which mark points of evolutionary divergence. These shell nodes often represent horizontal gene transfer (HGT) events and are commonly associated with alternate evolutionary pathways. We noted node differences between shell partitions that were consistent with virulence, metabolism, and defense systems, warranting further investigation.

### Virulence factors and predicted pathogenicity across *Gardnerella* lineages

Colonization of the vaginal niche by *Gardnerella*, and its transition to a pathogenic state, depend on mucin degradation, adherence to the vaginal epithelium, and the formation of structurally stable biofilms^13,55^. Canonical *Gardnerella* virulence factors implicated in these processes include exported sialidases and vaginolysin^56^, a cholesterol-dependent cytolysin. Among *Gardnerella*-associated sialidase isoforms, NanH2 and NanH3 account for the majority of extracellular sialidase activity, whereas NanH1 lacks a signal peptide and is therefore predicted to function intracellularly for sugar catabolism^57^. Consistent with this role, *nanH1*^58^ is embedded within a conserved operon across genomes (Extended Figure 5e). In contrast, *nanH2* and *nanH4*^59^ are frequently located in proximity to this operon but are associated with mobile genetic elements, suggesting more recent paralogous diversification and horizontal acquisition. Notably, *nanH3* is uniquely not operon-associated; instead, it is found as a solitary gene within defense-associated genomic regions, often adjacent to prophage elements, consistent with a distinct evolutionary trajectory and potential phage-linked mobility. Within our pangenome dataset, we identified lineage-specific nodes encoding vaginolysin, which contributes to host interaction by binding to and lysing epithelial cells of the vaginal mucosa^55^, and multiple enzymes involved in the stepwise degradation of mucin-associated glycans (Fig. 4d-e, Extended Data Fig. 5, Supplementary Table 5). These include secreted sialidases and additional carbohydrate-active enzymes, such as L-fucosidase^60–62^ and galactosidase^27^ that facilitate mucin catabolism when exported^63^.

A Type II α-amylase-pullulanase protein was well conserved (present in 100% of *Gardnerella* genomes), in agreement with previous reports^64^ (Fig. 4d). Comparisons between *Gardnerella* Set A and Set B showed that the four sialidase genes (*nanH1*-*nanH4*) are significantly enriched in Set A genomes, while vaginolysin (*vly*) is significantly enriched in Set B genomes (Fig. 4e, Extended Data Fig. 5), highlighting lineage-dependent differences in metabolic capacity and the distribution of putative virulence-associated genes across *Gardnerella*.

### Predicted metabolic capabilities across *Gardnerella* lineages

Given the observed difference in metabolism-related nodes, we bioinformatically predicted the biosynthesis of amino acids and the catabolism of small carbon sources across our *Gardnerella* genome dataset (Fig. 5a, Fig. 5f, Supplementary Table 6). The biosynthesis of glutamine, glycine, and proline was well conserved (>99% of genomes) while a high proportion of genomes (89%) harbored the capacity for cysteine biosynthesis (Fig. 5a). Asparagine biosynthesis was enriched for overall for Set B versus Set A, but within Set A, 100% of the *Gr* strains carried the genetic pathway. Within Set B, asparagine biosynthesis was enriched for *Gg* (although depleted for *Ggi*) and the *Gll* lineage alone amongst other *Gl* (Fig. 5a, b). Methionine biosynthesis is also enriched throughout Set B, but absent in all of Set A. Within Set B methionine biosynthesis is notably lacking for strains of *Gb* and *Ggg* (Fig. 5a, Fig. 5d). Threonine biosynthesis is much more sporadically throughout Set A as compared to Set B (Fig. 5a, Fig. 5e). Additionally, threonine synthesis is more likely to be found in *Gv* compared to other Set A (Fig. 5a, Fig. 5e). Although cysteine biosynthesis is well conserved this pathway is entirely absent in *Gh* genomes (Fig. 5a) and depleted in *Gps* as compared to other *Gp* genomes (Fig. 5a, Fig. 5e).

**Figure 5:**
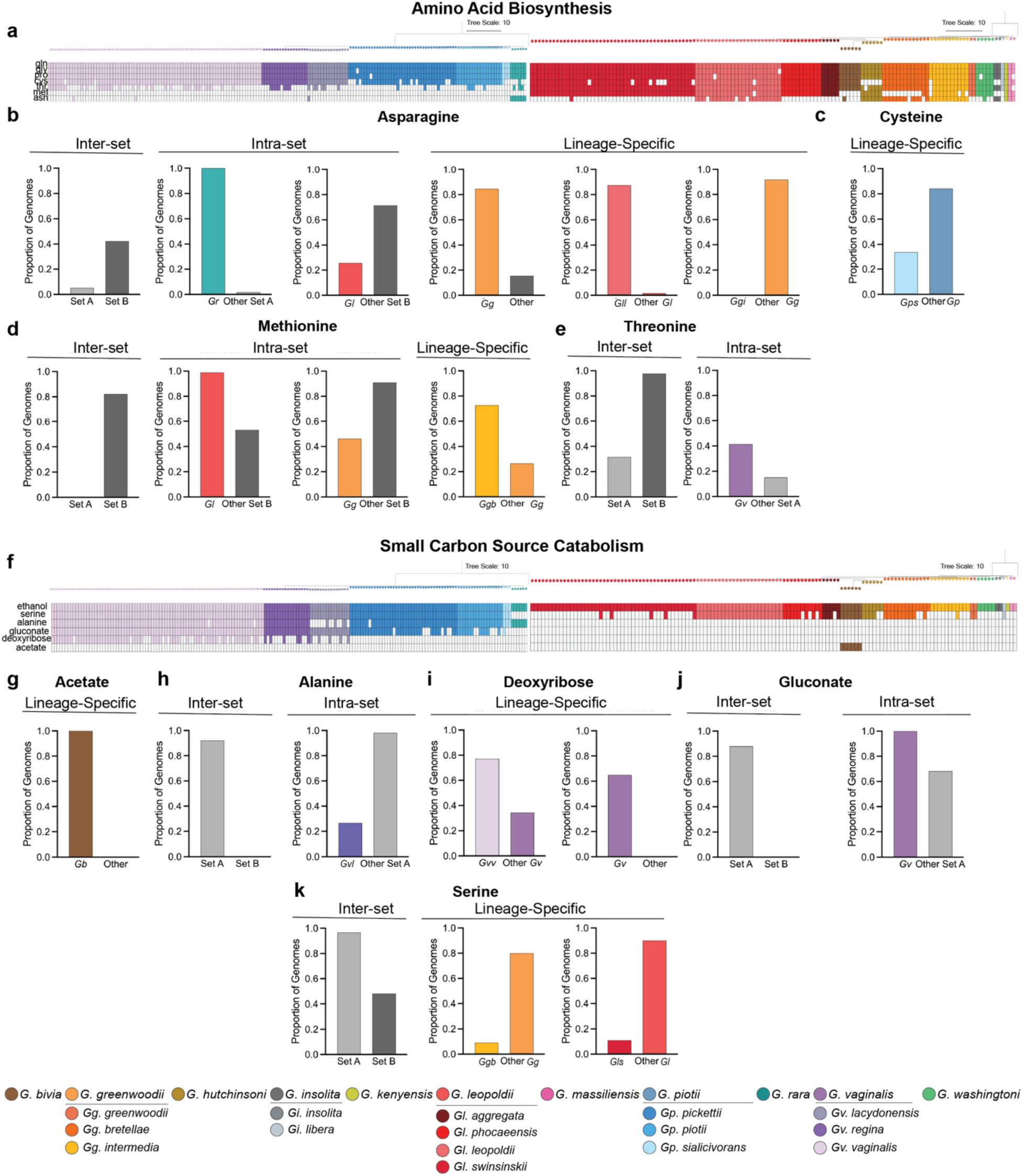
Predictions for metabolic capabilities across *Gardnerella* lineages. Presence-absence map for GapMind-predicted pathways to **a,** synthesize amino acids (AA) or **f,** catabolize small carbon sources (SCS) where each row represents a *Gardnerella* genome. Genomes are ordered by kSNP4 phylogenies of Set A and Set B. Presence by proportion comparison of genomes for the **b-e**, biosynthesis of a subset of amino acids or **g-k**, catabolism of a subset of small carbon sources. Colors represent each lineage.

Genes for the catabolism of ethanol appear to be a well-conserved feature of *Gardnerella* genomes (Fig. 5f). Genomes across Set A were additionally significantly enriched for the catabolism of alanine, gluconate, and serine (Fig. 5f, Fig. 5h, Fig. 5j, Fig. 5k). Notably, lineage-specific associations within Set A were observed for the catabolism of each of these small carbon sources. While gluconate catabolism is significantly enriched in *Gv*, alanine catabolism is significantly depleted in *Gvl* as compared to other Set A members (Fig. 5f, Fig. 5h, Fig. 5j). Serine catabolism was completely absent in *Gr* and (Fig. 5f) and was significantly depleted in *Ggb* and *Gls* compared to other lineages within their respective species (Fig. 5f, Fig. 5k). Interestingly, the catabolism of acetate and deoxyribose was lineage-specific and unique to *Gb* and *Gv*, respectively (Fig. 5f). Deoxyribose catabolism was enriched in the *Gvv* compared to other *Gv* subspecies (Fig. 5f, Fig. 5i). Our results indicate that Set A and Set B genomes are distinguished by their relative enrichment of pathways for catabolism of small carbon sources or synthesis of distinct amino acids with lineage-specific associations.

### Phage and antiphage defense mechanisms across *Gardnerella* genomes

In our pangenome data set, we noted that multiple nodes were related to phage-associated proteins that encompassed two distinct *Siphoviridae* family *Gardnerella* prophage, including the previously described vB_Gva_AB1^51,65^ and a novel phage we have tentatively designated as “pGamma” (Extended Data Fig. 5, Supplementary Table 7). This newly recognized phage is from a distinct family from vB_Gva_AB1 within the Class *Caudoviricetes* in whole genome comparisons^66^. vB_Gva_AB1 is sometimes reported as several different prophages^46,51^, the discrete synteny within our PPanGGOLiN pangenome suggests that these variants represent at least one discrete taxon (Extended Data Fig. 5). It is notable that some strains appear to have remnant phage genes, although these appear to be non-syntenic and scattered throughout the genome (Supplementary Table 5). While both vB_Gva_AB1 and pGamma were identified in Set A and Set B genomes, vB_Gva_AB1 was significantly enriched in the Set B species *G. leopoldii* (Bonferroni adj. p < 0.01, Fisher’s Exact test) and *G. greenwoodii* (Bonferroni adj. p = 0.4, Fisher’s Exact test) (Extended Data Fig. 6).

To counteract phages, bacteria evolved an array of defense systems^67^, with the most widespread and well-characterized being restriction-modification (RM)^68,69^ and CRISPR-Cas mechanisms^70^. Across our *Gardnerella* genome dataset we identified the presence of sixty different systems, with an average of twelve defenses per genome (Fig. 6a-b, Extended Data Fig. 7, Supplementary Table 8). Of the defense systems with described mechanisms, dXTPase systems^71^ were well conserved (99% genomes). Type I RM systems were prevalent (56% of genomes) and did not differ significantly between *Gardnerella* sets. Set A genomes were significantly enriched for some Abi proteins (AbiD^72^ and AbiL^73^), CBASS defenses^74,75^, and DRT mechanisms^76,77^, while Set B showed enrichment for a larger number of defenses, including those with Abi-mechanisms (AbiE^69^, Gabija^75^, and Lamassu^75,78^), CRISPR-Cas (CRISPR arrays and Cas Type IE proteins), and Type II RM defenses (Extended Data Fig. 8). Across *Gardnerella* lineages, *Gls* showed an enrichment for DNA-modification system (DMS) proteins, while *Gvv* showed an enrichment for AbiL and DRT defenses (Extended Data Fig. 8).

**Figure 6:**
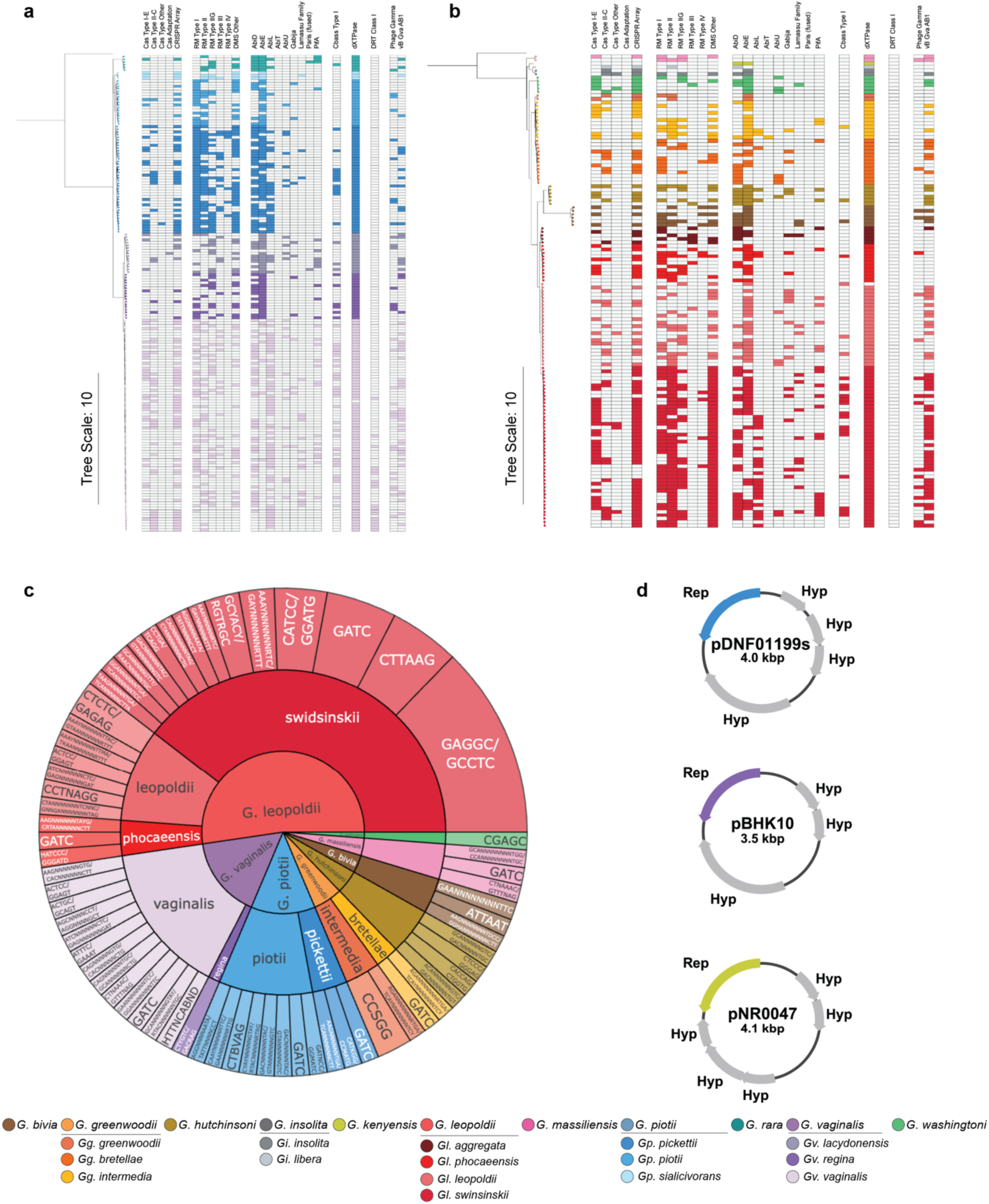
Defense systems, methylation motifs, and plasmid architectures across *Gardnerella* lineages. **a-d,** Presence-absence maps for PADLOC-predicted defense mechanisms. Genomes are ordered by kSNP4 phylogenies of **a,** Set A and **b,** Set B. **c,** Sunburst plot shows the sequence of each methyl-modified motif detected within each *Gardnerella* lineage. Size of section represents prevalence within each lineage. **d,** Schematics depict plasmid annotations for pDNF01199S-01, p0756_546_1_1_BHK10-01, and pNR047-01. Full sequences for the plasmids are available in Supplementary Table 10.

Although defense mechanisms evolved primarily to protect against bacteriophages, their activity extends against other horizontally transferred genetic elements, as well as plasmid vectors used during *in vitro* transformation. RM defenses, which are dependent upon detection and destruction of inappropriately methylated foreign DNA, are one of the most prevalent across bacteria. Further, it is notable that approaches to overcome these RM systems during genetic engineering have been shown to significantly improve transformation efficiency across multiple bacterial species^79–81^. Thus, we sought to characterize RM defenses across a subset of our *Gardnerella* genome dataset. We obtained and analyzed methylomes for 42 *Gardnerella* isolates across thirteen lineages within the Fredricks isolate collection (Supplementary Table 2). A total of 39 methylated motifs were detected and assigned to Type I (25 motifs), Type II (11 motifs), or Type III (3 motifs) RM systems (Fig. 6c, Supplementary Table 9).

### Identification of a native *Gardnerella* cryptic plasmid

There is renewed interest in developing genetic tools for engineering *Gardnerell*a species^82,83^. Because *Gardnerella* was long thought to lack mobile genetic elements, such as native plasmids, most efforts have focused on adapting non-*Gardnerella* genetic tools for use in this genus. Recent studies have been extensive and show promising results, achieving targeted chromosomal integrations and deletions using *E. coli*-based suicide vectors^82^, as well as reproducible transformation with a *Bifidobacterium-E. coli* shuttle vector^83^. During genome sequencing of the Fredricks clinical isolate collection, we observed an independent circular contig, with high coverage, forming a second replicon within the otherwise closed genome sequence of *Gppk* strain DNF01199S (Fig. 6d, Supplementary Table 10). Annotation of this contig showed three coding sequences for hypothetical proteins and a single putative replication initiation protein (Supplementary Table 11). Subsequent plasmid isolation and validation confirmed the episomal nature of this contig, marking pDNF01199S-01 as the first cryptic plasmid identified within the *Gardnerella* genus.

Motivated by this discovery, we assessed whether additional cryptic plasmids were present in our *Gardnerella* pangenome. We reasoned that the current limited knowledge of such plasmids reflects their rarity. As such, we focused on the cloud partition of the accessory genome, which highlights strain-specific genetic factors and collectively comprises 47.23% of the *Gardnerella* pangenome. Using structural sequence insights from pDNF01199S-01, we identified two additional putative plasmid sequences within the cloud partition: one in *Gvv* strain 0756_546_1_1_BHK10^84^ (p0756_546_1_1_BHK10-01), which was isolated from cervicovaginal mucus in South Africa, and another in *Gk* strain NR047^23^ (pNR047-01), isolated from a vaginal swab in Nairobi, Kenya. All three plasmids are approximately 4kb in size, contain coding sequences for putative hypothetical proteins, and share a highly similar replication initiation protein (84.48%-95.26% similar), suggesting a potentially conserved replication mechanism for cryptic plasmids within the *Gardnerella* genus (Fig. 6d, Supplementary Tables 10-11). Given that pDNF01199S-01 replicates stably in *Gardnerella*, we next sought to engineer a *Gardnerella-E. coli* shuttle vector suitable for transformation of additional *Gardnerella* isolates. Analysis of our pangenome revealed that 22% of *Gardnerella* genomes encode *tetM*, including *Gppk* DNF01199S, consistent with previous reports on variable tetracycline resistance phenotypes across *Gardnerella* strains^85^. We therefore assessed the growth of both *tetM*+ and *tetM*- isolates in the presence of supplemented tetracycline. As predicted, *tetM*+ isolates were resistant to tetracycline, whereas *tetM*- isolates were susceptible, confirming that the *tetM* cassette confers tetracycline resistance *in vitro*. This supports the use of *tetM* as a selectable marker for this vector. To enable the propagation in *E. coli* strains, we combined pDNF01199S-01, the *tetM* cloned from DNF01199S, and an *E. coli* minicircle plasmid backbone (pMC) into a single <u>p</u>arental <u>p</u>lasmid (herein designated pG01-pp) (Extended Data Fig. 9a-c). As previously reported^86^, we induced and purified the pDNF01199S-tetM-MC <u>m</u>ini<u>c</u>ircle vector (pG01-MC). We initially attempted to transform this minicircle into the *tetM*-strain, ATCC 14018^T^ (*Gvv*), but no transformants were recovered (Extended Data Fig. 9d).

The strain ATCC 14018^T^ encodes a restriction-modification (RM) system which modifies the “GGCC” motif^84^ and has been shown to limit transformation efficiency^82^. Thus, we used commercially available *HaeIII* methyltransferase (MTase) to methyl-modify these sites in pG01-MC prior to transformation. Electroporation with methylated pG01-MC successfully transformed *Gvv* ATCC 14018^T^, but at a very low efficiency (two transformants per µg of DNA). Further optimization of the protocol increased efficiency to 685 transformants per µg of DNA by transforming with 0.5 µg total *HaeIII*-treated vector on competent cells grown with 100 mM NaCl to an OD_600_ of 0.32 and washed with 10% glycerol (Extended Data Fig. 9d). A detailed, step-by-step version of the optimized protocol is available online at (https://www.protocols.io/private/1A47BAD2F1AC11F09AA30A58A9FEAC02).

Collectively, these results show that pG01-pp is the first replicative shuttle vector derived from a native *Gardnerella* cryptic plasmid. Together with recent work by the Dillard and Hill groups^82,83^, these efforts should establish a robust platform for more mechanistic investigations of *Gardnerella* biology and pathogenesis.

## Discussion

Taxonomic inconsistencies and fluctuating nomenclature have made it inherently challenging to delineate the roles of *Gardnerella* lineages in health and disease. In this study, we integrate whole-genome analyses to establish a refined and internally consistent taxonomic framework for the *Gardnerella* genus. Using a curated set of 313 distinct, high-quality genomes, we define clear lineage boundaries and propose an updated *Gardnerella* nomenclature that reflects genome-scale divergence while preserving continuity with existing naming conventions. Although nomenclatural revisions, particularly for well-established organisms such as ‘*G. vaginalis,’* (updated here to *G. vaginalis* subsp. *vaginalis*), can often be contentious, applying established genomic criteria for species and subspecies delineation is warranted to prevent further taxonomic ambiguity and this misinterpretation of functional studies.

While single-locus markers such as *cpn60*, which are widely used for *Gardnerella* typing, can distinguish between *Gardnerella* species, they lack sufficient resolution for subspecies assignments. Accurate subspecies assignments require whole-genome approaches including phylogenomic inference and average nucleotide identity or digital DNA-DNA comparisons. As additional *Gardnerella* lineages are isolated, whole-genome sequencing efforts should allow for continuous integration into this framework, supporting a stable and extensible taxonomy.

Our genome-based analyses also intersect with a broader, long-standing debate regarding the placement of *Gardnerella* within the family *Bifidobacteriaceae*. Although, marker gene frameworks such as the GTDB place *Gardnerella* within *Bifidobacterium*, the two groups differ markedly in genome size, G+C content, metabolic repertoire, virulence-associated gene content, mobile genetic element composition, and ecological specialization. Notably, *Gardnerella* genomes are substantially smaller and lack many biosynthetic and metabolic pathways retained in larger *Bifidobacterium* genomes. Such genome reduction is widely interpreted as a hallmark of specialization, reflecting adaptation to host-associated niches where broad metabolic versatility is unnecessary. In this context, reduced genome size represents ecological specialization rather than diminished biological significance.

Together, these genome-scale differences suggest that conserved marker phylogeny alone may be insufficient to define biologically meaningful genus boundaries within this family. A parsimonious interpretation is that *Bifidobacterium*, as currently circumscribed, encompasses multiple genomically and ecologically distinct genera. While a formal genus-level revision of *Bifidobacteriaceae* lies beyond the scope of this study, our data provide additional genome-scale evidence supporting the continued recognition of *Gardnerella* as a distinct genus and contribute to an evidence base that may inform future ICNP-compliant taxonomic restructuring. With this refined taxonomic and nomenclature framework, we probed for genetic features and putative operons associated with each lineage by applying a syntenic pangenome approach. These analyses linked *Gardnerella* lineages to distinct repertoires of virulence-associated factors, metabolic pathways, and genetic defense systems, highlighting functional diversity across the genus. Notably, most *Gardnerella* gene content (84%) is species- or strain-variable with many differences related to virulence-associated functions, metabolism, and genetic defense.

*Gardnerella* Set A genomes (*Gr, Gp,* and *Gv*) were enriched for sialidases and exhibited increased potential for the catabolism of small carbon sources. We further noted lineage-specific enrichment of established virulence-associated factors, including exported sialidases (*nanH2* in *Gr* and *Gppt*; *nanH3* in *Gppk* and *Gppt*) and vaginolysin (enriched in *Gvv, Gvl, Gps* and *Gppk*). The restricted distribution of these factors suggests functional specialization among lineages that may contribute to mucin degradation and epithelial barrier disruption within the vaginal microbiome. In contrast, Set B genomes (*Gb, Gc, Gg, Gi, Gl, Gk, Gm,* and *Gw*) were enriched in putative virulence-associated proteins (e.g. predicted surface-associated proteins, secretion system components and factors implicated in host interaction), exhibited greater capacity for amino acid biosynthesis, showed a higher propensity to harbor the vG_gva_AB1 prophage, and encoded a broader repertoire of genetic defense mechanisms. Collectively, these features are consistent with distinct colonization strategies among Set B *Gardnerella* lineages, including enhanced persistence on epithelial surfaces, flexibility under nutrient-limited or variable resource conditions, and adaptation to competitive, phage-rich, polymicrobial niches.

Mechanistic studies to assess the roles of *Gardnerella* genetic factors, including those identified in this study, have been hindered by limited genetic tools available for *Gardnerella*. In fact, it has previously been reported that *Gardnerella* species lack mobile genetic elements, such as plasmids^30^. However, our extensive genome sequencing efforts reveal the first *Gardnerella* cryptic plasmid naturally occurring within a single cultivated strain, along with additional homologous sequences in publicly available genomes. Through the identification of this cryptic plasmid (pDNF01199S), we developed a replicative *Gardnerella-E. coli* shuttle vector (pG01-pp). Coupled with recent advances in *Gardnerella* species engineering^82,83^, this discovery of extrachromosomal and self-replicative genetic elements within the *Gardnerella* genus should facilitate the development of advanced genetic tools. Combined, these foundational studies seek to advance genetic tools for mechanistic studies of *Gardnerella* species. Collectively, overcoming genetic intractability within *Gardnerella* will greatly advance our ability to investigate the role of this genus in health and disease states.

Although these interpretations are based on comparative genomic potential and do not directly infer *in vivo* activity or clinical outcomes, taken together they support the view of *Gardnerella* as an ecologically diverse genus composed of lineages with distinct metabolic capacities, host-interaction mechanisms, and mobile genetic repertoires. Different *Gardnerella* seem to occupy overlapping but not identical niches within their human hosts. Some lineages may be better adapted to persistence and competitive growth in the vaginal microbiome and urinary tract, whereas others may be more consistent with transient or context-dependent colonization.

## Methods

### Study design and participants

Vaginal bacterial isolates were collected from human study participants both with and without BV in Seattle, Washington, USA^22,87^ or from the Ravel isolate collection, harboring strains from multiple geographic locations, including USA and Africa (Supplementary Table 2). Bacterial isolates consistent with *Gardnerella* morphotypes on Gram stain were subjected to 16S rRNA and/or *cpn60* gene amplicon sequencing to confirm phylogenetic placement in the *Gardnerella* genus^24^. Phylogenetically diverse *Gardnerella* species were selected for whole-genome sequencing. High molecular weight DNA was extracted from each isolate using the MasterPure DNA purification kit (Lucigen; Middleton, WI), with two phenol/chloroform cleanups prior to DNA precipitation where needed. DNA extraction was quantified on a TapeStation 2200 instrument run with a Genomic DNA tape (Agilent Technologies; Santa Clara, CA).

### Genome sequencing and assembly

#### Sequencing on the Pacific Biosciences SeqI/II Platforms

A total of forty-two strains from the Fredricks collection, and seventeen from the Ravel collection, were selected for single-molecule real-time (SMRT) long-read sequencing using PacBio (SeqI or SeqII, Pacific Biosciences; Menlo Park, CA) at the Fred Hutch Cancer Center or the Genomic Resource Center of the University Maryland School of Medicine. Sequencing libraries were prepared using the SMRTBell Template Prep Kit 1.0/2.0 and were selected for a 3-5kb size cutoff using a BluePippin (Sage Science; Beverly, MA). Sequencing reads were processed via Microbial Assembly software within the Pacific Biosciences’ SMRTAnalysis pipeline (v.9.0.0.92188).

#### Sequencing on the Illumina HiSeq 4000 Platform

332 isolates were selected for Illumina sequencing on the HiSeq4000 platform. Short read libraries were prepared using the KAPA HyperPlus Kit (Kapa Biosystems/Roche; Willmington, MA) with KAPA Single-Indexed Adapter Kit Set B. 35μL of genomic DNA was used as input, and libraries were prepared following the manufacturer’s protocol with modifications based on concentration. For samples with 0.5 or 0.2 ng input DNA, the fragmentation enzyme was diluted 1:2 or 1:5 with water. All samples were fragmented at 37°C for 5 minutes. Adapter concentrations varied according to the input, and the adapter ligation was carried out overnight at 4°C for all samples. The post-ligation cleanup was performed with 0.8x Ampure XP beads (Beckman Coulter; Indianapolis IN) and 20 μL of sample was used in library amplification. Post-amplification cleanup was performed with 1x Ampure XP beads; libraries with remaining adapter dimer peaks were cleaned a second time. The final elution was in 25 μL of nuclease-free water. Libraries were run on a TapeStation instrument with a D1000 tape (Agilent Technologies; Santa Clara, CA) to assess quality and concentration. Libraries were sequenced on an Illumina HiSeq 4000 instrument using the 150 bp paired-end protocol. Reads were assembled using Unicycler.

### Publicly available genomes from the National Center for Biotechnology Information (NCBI)

We used publicly available genomes both to provide a direct reference for previous studies and to supplement our own sequencing efforts. We searched GenBank^88^ assembly databases for the genus *Gardnerella* and family *Bifidobacteraceae* (without genus assignment). Using the Type (Strain) Genome Server (TYGS)^89^ to confirm the genus of these genomes, we discovered that several of the unassigned *Bifidobacteraceae* bacteria in NCBI were *Gardnerella* and included them in our analysis. *Bifidobacteraceae* (without genus assignment) were included if they were collected as part of the same NCBI BioProject which added 20 *Gardnerella* that were not typed further than the *Bifidobacteraceae* family. Additionally, we included three sequences (350_GVAG, 476_GVAG, and 842_GVAG) that were initially assigned to *Gardnerella* but were later renamed to *Alloscardovia* to verify this placement. The initial number of potential genomes was n = 649. We removed genomes that were assigned outside of *Gardnerella*. Draft genomes with more than 250 contigs, modified in the laboratory, and those with truncated or missing cpn60 sequences were removed due to inherent unreliability.

*Bifidobacterium* strains were collected and curated in much the same manner as the *Gardnerella* strains. A total of 98 strains from all known species of the *Bifidobacterium* genus and the closely related scardovial genera (e.g., *Alloscardovia, Aeriscardovia*, *Bombiscardovia*, *Pseudoscardovia*, and *Scardovia*) were downloaded from NCBI. Whenever possible, the type strains were collected. Sequences were assessed using CheckM, TYGS, and ANI via cutoffs similar to those used for the *Gardnerella* strains. *Bifidobacteriaceae* groups were assigned according to Alessandri *et al*^90^.

### Genome quality and comparison assessments

Genome completeness and contamination for all newly sequenced and publicly available *Gardnerella* genomes retrieved from NCBI were assessed using CheckM (v 1.0.18)^91^. All genomes were annotated via RAST (v 1.073)^92^. Genomes with contamination greater than 1.5% or completeness below 90% were excluded from downstream analyses. Redundant genome sequences were identified based on pairwise average nucleotide identity (ANI). Genomes sharing > 99.5% ANI with other genomes in the collection were considered duplicates and a single representative genome was retained for each duplicate group (Supplementary Table 1). Pairwise average nucleotide identity (ANI) metrics were calculated separately for Set A and Set B genomes using FastANI (v1.34)^93^ and visualized using the R packages ComplexHeatmap (v 2.27.0)^94^. Digital DNA-DNA hybridization (dDDH)^95^ values were calculated via the Genome-to-Genome Distance Calculator (v 3.0)^96^ and visualized in R (v 4.5.1 -- 2025-06-13 -- “Great Square Root”). Amino Acid Identity (AAI) values for *Bifidobacteriaceae* type strains were determined using pairwise comparisons of FastAAI (version 2)^44^.

### PPanGGOLiN *Gardnerella* pangenome

We used Partitioned PanGenome Graph of Linked Neighbors (PPanGGOLiN v 2.2.0)^52^ on our curated genome set to graph and cluster the genes from all strains, find regions of genomic plasticity and synteny^75^, and separate the pangenome gene clusters into core, shell, and cloud partitions, with clustering set to 50% coverage and 50% identity. EggNOG^96^, InterPro (accessed October 2025)^97^, and KEGG Mapper (v 5.2)^98^, were run separately on each genome to gather and confirm the CAZyme^99^, Pfam^100^, and KEGG (all accessed October 2025), designations of the gene clusters. COG categories^101^ were assigned based on the consensus of these designations. We added the annotation “Unclassified Metabolism” to capture gene clusters that are clearly related to metabolism, but the exact metabolite is not defined or could be of multiple types.

### Phylogenomic analyses

Barrnap was used to find and extract 16S rRNA sequences for submission to NCBI^98^. The sequences in the PPanGGOLiN for the persistent *cpn60* node were MUSCLE^99^-aligned to the same gene from *Bifidobacterium bifidum* (ATCC 29521) to generate maximum-likelihood distances, and a neighbor-joined dendrogram using the R package phangorn^100^. The resulting tree was visualized and annotated using iTOL (v 7.3)^101^. kSNP4 (v 4.1)^102^ was used to construct a reference-free whole-genome phylogenetic tree for *Gardnerella* Set A (kmer=17, FCK=0.167) and *Gardnerella* Set B (kmer=17, FCK=0.797) genomes, separately. *Bombiscardovia bombi* strain DSM19703 was included to root both dendrograms. The resulting parsimony trees were visualized in iTOL^101^.GapMind (accessed September 2025)^103–105^ metabolism predictions and PADLOC (v 2.0.0)^106,107^ defense mechanism predictions were annotated where appropriate. Bacteriophage detection was performed by VIBRANT (v. 1.2.1)^108^, PHageR (accessed Feb 2026)^109^, and the use of phage-annotated nodes to find syntenic modules of phages in PPanGGOLiN. These methods were combined as no one method found all bacteriophage. Phage sequences are mostly as found by VIBRANT, when VIBRANT failed to find the phage sequence they were determined from PPanGGOLiN. Phages were typed by aligning the large terminase sequences from each intact with MAFFT (v7.526) and then creating a maximum likelihood tree with IQTree2 (v. 2.3.6)^110^. Assessment of greater taxonomic grouping was determined using VICTOR (https://ggdc.dsmz.de/victor.php accessed March 2026)^66^.

### Isolation of pDNF01199S-01 plasmid

*Gardnerella piotii* subsp*. pickettii* strain DNF01199S was grown anaerobically in New York City III (NYCIII) liquid media in an AS-580 anaerobic chamber (Anaerobe Systems, Morgan Hill, CA) in a gas mixture of 5% CO_2_, 5% H_2_, and 90% N_2_ at 37°C. A QIAamp DNA Miniprep kit (Qiagen, Hilden, Germany) was used to isolate pDNF01199S-01 from a 5 mL culture of the bacteria. Isolated DNA was digested with *XhoI* (New England Biolabs, Ipswich, MA) and visualized by gel electrophoresis to confirm plasmid presence.

### *Gardnerella* tetracycline resistance testing

*Gardnerella* strains were grown anaerobically in an AS-580 anaerobic chamber (Anaerobe Systems, Morgan Hill, CA) in a gas mixture of 5% CO_2_, 5% H_2_, and 90% N_2_ at 37°C. Bacteria were recovered from frozen stocks on NYCIII agar plates and subcultured twice in 2 mL of NYCIII liquid media. Culture purity was verified by Gram stain. The final cultures were normalized to 0.5 McFarland turbidity using fresh NYCIII broth and 25 µL was spotted onto an NYCIII agar plate containing 16 µg/mL tetracycline and an antibiotic-free NYCIII agar plate. 25 µL of *Lactobacillus reuteri* CF48-3A was spotted onto each tetracycline plate as a tetracycline-resistant control^102^. Tetracycline resistance was determined according to Clinical and Laboratory Standards Institute for antimicrobial susceptibility testing protocols, and any growth on tetracycline NYCIII agar plates was considered resistant to tetracycline.

### pMC-pDNF01199S-*tetM* construction and minicircle preparation

pDNF01199S-01 and *tetM* were both cloned from gDNA isolated from strain DNF01199S. Gene fragments were assembled with pMC by Gibson assembly^103^. The parental construct was transformed into *E. coli* ZYCY10P3S2T (System Biosciences, Palo Alto, CA) by heat shocking according to the SBI Minicircle DNA Technology user manual. Transformant colonies were selected by growing on Luria-Burtani (LB) agar plates containing 50 µg/mL Kanamycin. A successfully transformed colony was chosen and stocked, then induction of minicircles was performed according to the SBI Minicircle DNA Technology user manual by growing the bacteria in induction media containing arabinose. pDNF01199S-*tetM* minicircle DNA was then purified from the induced *E. coli* using a Qiagen Plasmid Midi Kit (Qiagen, Hilden, Germany). The sequence of purified plasmid was confirmed against the construct sequence by whole-plasmid sequencing performed by Plasmidsaurus (Plasmidsaurus, Eugene, OR). “GGCC” sites on the plasmid were protected by incubating 1mg of the DNA with 1µL of *HaeIII* MTase (New England Biolabs, Ipswich, MA) at 37°C for 4hrs in the presence of 160 µM S-Adenosyl methionine (New England Biolabs, Ipswich, MA). Following enzyme treatment, DNA was purified with Monarch PCR & DNA Cleanup Kit (New England Biolabs, Ipswich, MA).

### Preparation of competent *Gardnerella* cells

*Gardnerella vaginalis* subsp. *vaginalis* ATCC 14018^T^ was grown anaerobically in an AS-580 anaerobic chamber (Anaerobe Systems, Morgan Hill, CA) in a gas mixture of 5% CO_2_, 5% H_2_, and 90% N_2_ at 37°C. Cells were recovered from frozen stocks on NYCIII agar plates and subcultured into NYCIII liquid media. 500 µL of culture was subcultured into 5 mL NYCIII media and grown overnight. 4mL of this culture was inoculated into 40mL of modified de Man-Rogosa-Sharpe media^104^ supplemented with 5 g/L glucose, 0.4 g/L L-cysteine HCl, 100 mL/L filter-sterilized horse serum, and 100 mM NaCl when stated. Once the target OD_600_ was reached, cells were harvested by centrifugation at 4°C at 4,052 x *g* for 10 minutes. Cells were washed in one of the following three ways: 1) Twice in 25 mL ice-cold maltose citrate buffer (0.2625 g/L citric acid and 0.5 M maltose at pH 5.8), 2) Twice in 25 mL ice-cold, autoclaved, 10% glycerol, or 3) Once in 25 mL ice-cold autoclaved water from a Milli-Q IQ7000 pure water system (Millipore Sigma, St. Louis, MO) followed by a second wash in 25 mL ice-cold maltose citrate buffer. Washed cells were then resuspended in 200 µL of maltose citrate buffer, mixed with 200 µL 80% glycerol, and stored at -80°C in aliquots.

### *Gardnerella* electrotransformation and verification

Competent cells were mixed with plasmid DNA on ice and transferred into pre-chilled 0.1 cm gap Gene Pulser/MicroPulser Electroporation Cuvettes (Bio-Rad, Hercules, CA). Electroporation was then performed on a Gene Pulser Xcell (Bio-Rad, Hercules, CA) at 25 µF, 200 Ω, and 2,000 V. Immediately following electroporation, 950 µL of 37°C pre-warmed NYCIII broth was added to the cells and they were transferred to an AS-580 anaerobic chamber (Anaerobe Systems, Morgan Hill, CA) in a gas mixture of 5% CO_2_, 5% H_2_, and 90% N_2_ to recover at 37°C for 1 hour. 100 µL – 200 µL of transformed cells were then plated on NYCIII agar plates with and without 16 µg/mL tetracycline. Plates were incubated anaerobically at 37°C for 48 hours and transformed colonies on the tetracycline plates were counted to determine transformation efficiency. After the first successful transformation, transformant colonies were grown in 5 mL NYCIII media with 16 µg/mL tetracycline and pDNF01199S-*tetM* was recovered from them using a QIAamp DNA Miniprep kit (Qiagen, Hilden, Germany). The sequence of the purified plasmid was confirmed by sequencing performed by Plasmidsaurus (Plasmidsaurus, Eugene, OR). Following transformation of *Gardnerella vaginalis* subspecies *regina* strain DNF00354 with pDNF01199S-*tetM*, the transformation reaction was plated onto NYCIII agar plates with and without 16 µg/mL tetracycline. A small amount of growth from a transformant colony on the tetracycline plate or lawn growth from the control plate was resuspended in sterile water and colony PCR was performed to amplify and detect the *tetM* gene.

One microliter of resuspended bacteria was added to a reaction mix of 12.5 µL GoTaq Green master mix (Promega, Madison, WI), 0.5 µL of each primer, and 6.5 µL H_2_O. PCR was carried out for 28 cycles with an annealing temperature of 60°C and an extension time of 2.5 min. A detailed version of this protocol is available online at: https://www.protocols.io/private/1A47BAD2F1AC11F09AA30A58A9FEAC02.

Primer sequences were as follows (5’ to 3’):

Forward: CGCGACCAAATATTGGTACATTATTACAGCTATTTTGTAATCACGTACTC

Reverse: GAGACTGTCGTATTGATGTCACGGGCTTAATAGGTCTTATGTCGG

### Protologues

Description of *Gardnerella vaginalis* subsp. *vaginalis* Gardner and Dukes 1955 (Approved Lists 1980) Bouzek *et al*. comb. nov.

Basonym: *Haemophilus vaginalis* Gardner and Dukes 1955 (Approved Lists 1980)^9^. Basonym:

*Haemophilus vaginalis* Gardner and Dukes 1955 (Approved Lists 1980)^9^.

Earlier homotypic synonym: *Gardnerella vaginalis* (Gardner and Dukes 1955) Greenwood and Pickett 1980^114^. Earlier homotypic synonym: *Gardnerella vaginalis* (Gardner and Dukes 1955) Greenwood and Pickett 1980^111^.

*Gardnerella vaginalis* subsp. *vaginalis* (va.gi.na’lis. L. fem. n. *vagina*, sheath, vagina; L. masc. adj. suff. -*alis,* suffix denoting pertaining to; N.L. fem. adj. *vaginalis*, pertaining to vagina, of the vagina).

The description of this subspecies remains as given to *Gardnerella vaginalis* by Gardner and Dukes 1955^9^ and Greenwood and Pickett 1980^111^ with the following additional information. Curated genomes of 79 *Gardnerella vaginalis* subsp. *vaginalis* cultivated isolates were used in this analysis, isolated from the vagina, endometrium, cervix, catheter urine, and placenta of humans from Bangladesh, Belgium, China, France, Russia, the United Kingdom, the United States, and Zambia (Supplementary Table 2). The mean genome length of *Gardnerella vaginalis* subsp. *vaginalis* is 1.68 Mb [range1.58 – 1.78] and the median G+C content is 41.26% [range 40.99% – 41.82%]. The type strain for *Gardnerella vaginalis* subsp. *vaginalis* remains 594^T^ of Gardner and Dukes (=ATCC 14018^T^=DSM 4944^T^=CCUG 3717^T^=JCM 11026^T^=NCTC 10915^T^=NCTC 10287^T^).

Strains used in previous classifying publications: ATCC14019, ATCC49145, DNF01149, FDAARGOS296, FDAARGOS568, GH015, GH021, HMP9231, JCM11026, JCP7276, JCP7672, NR001, NR037, NR038, NR039, UGent09.01, UGent09.07, UGent25.49, UMB0032A, UMB0032B, UMB0061, UMB0233, UMB0298, UMB0386, UMB0768, UMB0770, UMB0775, WP023, 287, 377, 653, 1492, 3549624, 0284V, 0286E, 0288E, 1-010-6, 1-014-4, 104-V2-10, 1388E, 18-4, 203-8, 23-12, 302000480-V1-10, 315-A, 7571-2, 772-1, and 825-1.

Description of *Gardnerella vaginalis* subsp. *lacydonensis* (Allini Ntiguemassa *et al.* 2026) Bouzek *et al.* comb. nov.

Basonym: *Gardnerella lacydonensis* Allini Ntiguemassa *et al*. 2026^49^ Basonym: *Gardnerella lacydonensis* Allini Ntiguemassa *et al*. 2026^49^

*Gardnerella vaginalis* subsp. *lacydonensis* (la.cy.do.nen’sis. N.L. fem. adj. *lacydonensis*, from Lacydon, the name of the ancient port of Marseille, the French city where the strain was first described).

The description of *Gardnerella vaginalis* subsp. *lacydonensis* remains as given to *Gardnerella lacydonensis* by Allini Ntiguemassa *et al*.^49^ with the following additional information. Our collection of curated genomes contains 15 *Gardnerella vaginalis* subsp. *lacydonensis* cultivated strains, isolated from the vagina, endometrium, and ectocervical mucosa of humans from France, Kenya, South Africa, the United States, and Zambia (Supplementary Table 2). The median genome length of these isolates is 1.66 Mb [range 1.62 – 1.72] and the median G+C content is 41.29% [range 41.11% – 41.42%]. The type strain of *Gardnerella vaginalis* subsp. *lacydonensis* remains Marseille-Q9181^T^ (=CSUR Q9181^T^=CECT 31121^T^).

Description of *Gardnerella vaginalis* subsp. *regina* subsp. nov.

*Gardnerella vaginalis* subsp. *regina* (re.gi’na. L. fem. n. *regina*, the queen, referring to its placement in the first validly published species of the *Gardnerella* genus).

We analyzed 17 curated genomes from cultivated strains of *Gardnerella vaginalis* subsp. *regina*, isolated from the vagina of humans from Bangladesh, France, the United States, and Zambia (Supplementary Table 2). The median genome length of the *Gardnerella vaginalis* subsp. *regina* is 1.63 Mb [range 1.57 – 1.67], and the median G+C content is 41.41% [range 41.25% – 41.62%]. The isolate is a facultative anaerobe and has gram variable cells. Growth develops on Brucella agar in anaerobic conditions at 37°C after one to two days. In a genome comparison of *Gardnerella vaginalis* subsp. *regina* strain DNF00622A^T^ with validly published *Gardnerella* type strains and the type strains proposed in this study, the highest ANI values correspond with its sister subspecies *Gardnerella vaginalis* subsp. *vaginalis* ATCC14018^T^ (96.82%) and *Gardnerella vaginalis* subsp. *lacydonensis* Marseille-Q9181^T^ (96.17%), followed by *Gardnerella piotii* subsp. *piotii* UGent 18.01^T^ (89.24%) (Figure 2). Within the genome, 83 CAZymes were identified, including α-mannosidase (GH125), β-galactosidase (GH2), β-mannosidase (GH130), Class II α-mannosidase (GH38), β-N-acetylglucosaminidase (GH20), α-L-fucosidase (GH29), and sialidase (GH33)^112^. KEGG analysis identified a total of 97 transporters, 13 secretion genes, and 365 enzymes^112^. Metabolism predictions using GapMind^103–105^ include biosynthesis of cysteine, glutamine, glycine, and proline and catabolism of ethanol, gluconate, and serine (Supplementary Table 6). The type strain for *Gardnerella vaginalis* subsp. *regina* is DNF00622A^T^, isolated from the vaginal fluid of a woman positive for bacterial vaginosis in Seattle, Washington, USA. Accessions for the 16S rRNA gene sequence and whole genome sequence for DNF00622A^T^ are PZ013896 and CP058395.

Strains used in previous classifying publications: 1-004-3, 819-1, and 819-3.

Description of *Gardnerella piotii* subsp. *piotii* (Vaneechoutte *et al*. 2019) Bouzek *et al*. comb. nov.

Basonym: *Gardnerella piotii* Vaneechoutte *et al*. 2019^32^

*Gardnerella piotii* (pi.o’ti.i. N.L. gen. masc. n. *piotii*, of Piot, named after Peter Piot, a Belgian microbiologist who first attempted to distinguish different groups within *G. vaginalis*).

The description of *Gardnerella piotii* subsp. *piotii* remains as given to *Gardnerella piotii* by Vaneechoutte *et al*.^32^ with the following additional information. There are 17 genomes of cultivated *Gardnerella piotii* subsp. *piotii* in our curated collection, isolated from the vagina and endometrium of humans from Bangladesh, Belgium, China, France, and the United States (Supplementary Table 2). The median genome length of these isolates is 1.54 Mb [range 1.50 – 1.59] and the median G+C content is 42.43% [range 42.22% – 42.61%]. The *Gardnerella piotii* subsp. *piotii* type strain remains UGent 18.01^T^ (= LMG 30818^T^=CCUG 72427^T^).

Description of *Gardnerella piotii* subsp. *pickettii* (Sousa *et al*. 2023) Bouzek *et al*. comb. nov.

Basonym: *Gardnerella pickettii* Sousa *et al*. 2023^48^

*Gardnerella piotii* subsp. *pickettii* (pic.ket’ti.i. N.L. gen. masc. n. *pickettii*, of Pickett, named after M. John Pickett, an American microbiologist who proposed the transfer of *Haemophilus vaginalis* to the new genus *Gardnerella*).

The description of *Gardnerella piotii* subsp. *pickettii* remains as given to *Gardnerella pickettii* by Sousa *et al*.^48^ with the following additional information. We included the curated genomes of 40 cultivated isolates of *Gardnerella piotii* subsp. *pickettii* in our analysis, isolated from the vagina, endometrium, ectocervical mucosa, urine, bladder and rectum of humans from Bangladesh, Canada, China, Kenya, Portugal, South Africa, the United States, and Zambia (Supplementary Table 2). The median genome length of *Gardnerella piotii* subsp. *pickettii* is 1.56 Mb [range 1.51 – 1.68] and the median G+C content is 42.32% [range 42.00% - 42.74%]. The type strain of *Gardnerella piotii* subsp. *pickettii* remains c17Ua_112^T^ (=DSM 113414^T^=CCP 71^T^).

Description of *Gardnerella piotii* subsp. *sialicivorans* subsp. nov.

*Gardnerella piotii* subsp. *sialicivorans* (si.a.li.ci.vo’rans. N.L. neut. n. *acidum sialicum*, sialic acid; L. pres. part. *vorans*, devouring; N.L. fem. part. adj. *sialicivorans*, sialic acid devouring).

The subspecies *Gardnerella piotii* subsp. *sialicivorans* is named for the intact sialidase catabolism operon and one or more sialidase genes present in the genomes of *Gardnerella piotii*. Our collection of curated genomes includes three *Gardnerella piotii* subsp. *sialicivorans* cultivated strains, all isolated from the vagina of humans from the United States (Supplementary Table 2). The median genome length is 1.51 Mb [range 1.50 – 1.53], and the median G+C content is 42.48% [range 42.39% - 42.48%]. This isolate is a gram-positive coccus and produces pinpoint colonies (≤ 1 mm) on Brucella agar after three days of anaerobic incubation at 37°C. Colonies are grey, pleomorphic, and alpha-hemolytic. In a genome comparison of *Gardnerella piotii* subsp. *sialicivorans* strain C0013B2^T^ with validly published *Gardnerella* type strains and the type strains proposed in this study, the highest ANI values correspond with members of its sister subspecies *Gardnerella piotii* subsp. *pickettii* c17Ua_112^T^ (95.68%) and *Gardnerella piotii* subsp. *piotii* UGent 18.01^T^ (94.66%), followed by *Gardnerella rara* C0101A1^T^ (90.88%) (Figure 2). Within the genome, 76 CAZymes were identified, including sialidase (GH33) and Class II α-mannosidase (GH38)^112^. KEGG analysis identified a total of 79 transporters, 13 secretion genes, and 355 enzymes^112^. Metabolism predictions using GapMind^103–105^ include biosynthesis of glutamine, glycine, and proline, and catabolism of alanine, ethanol, and serine (Supplementary Table 6). The *Gardnerella piotii* subsp. *sialicivorans* type strain is C0013B2^T^, isolated from the vagina of an asymptomatic menstruating woman in Birmingham, Alabama, USA. Accession for the 16S rRNA gene sequence for C0013B2^T^ is PZ016854.

Strain used in previous classifying publications: 582.

Description of *Gardnerella rara* sp. nov.

*Gardnerella rara* (ra’ra. L. fem. adj. *rara*, rare, uncommon, referring to its rare genitourinary colonization of humans).

Our curated collection includes six genomes of *Gardnerella rara* from isolates cultivated from the vagina and endometrium of humans in Bangladesh, the United States, and Zambia (Supplementary Table 2). The median genome length of *Gardnerella rara* is 1.52 Mb [range 1.47 – 1.58] and the median G+C content is 43.23% [range 43.01% – 43.29%]. This isolate is a gram-positive bacillus and produces small colonies on Brucella agar after two days of anaerobic incubation at 37°C. Colonies are grey, round, and alpha-hemolytic. In a genome comparison of *Gardnerella rara* strain C0101A1^T^ with validly published *Gardnerella* type strains and the type strains proposed in this study, the highest ANI values correspond with *Gardnerella piotii* subsp. *piotii* UGent 18.01^T^ at 91.38% (Figure 2). Within the genome, 71 CAZymes were identified, including β-mannosidase (GH130), Class II α-mannosidase (GH38), and sialidase (GH33)^112^. KEGG analysis identified a total of 78 transporters, 13 secretion genes, and 344 enzymes^112^. Metabolism predictions using GapMind^103–105^ include biosynthesis of glutamine, glycine, and proline, and catabolism of alanine and ethanol (Supplementary Table 6). The type strain of *Gardnerella rara* type is C0101A1^T^, isolated from the vagina of a woman positive for *Chlamydia trachomatis* in Baltimore, Maryland, USA. Accession for the 16S rRNA gene sequence for C0101A1^T^ is PZ016855.

Strains used in previous classifying publications: 1401E, GED7760B, and HUZ601048-2.

Description of *Gardnerella leopoldii* subsp. *leopoldii* (Vaneechoutte *et al*. 2019) Bouzek *et al*. comb. nov.

Basonym: *Gardnerella leopoldii* Vaneechoutte *et al*. 2019^32^

*Gardnerella leopoldii* (le.o.pol’di.i. N.L. gen. masc. n. *leopoldii,* of Leopold, named after Sidney Leopold, who described the first strains of the species).

The description of this subspecies remains as given to *Gardnerella leopoldii* by Vaneechoutte *et al*.^32^ with the following additional information. There are 24 genomes of cultivated isolates of *Gardnerella leopoldii* subsp. *leopoldii* in our curated collection, isolated from the vagina and catheter urine of humans from Belgium, Canada, France, Kenya, and the United States (Supplementary Table 2). The median genome length is 1.53 Mb [range 1.37 – 1.67], and the median G+C content is 42.15% [range 42.07% - 42.34%]. The type strain of *Gardnerella leopoldii* subsp. *leopoldii* remains UGent 06.41^T^ (=LMG 30814^T^=CCUG 72425^T^).

Description of *Gardnerella leopoldii* subsp. *swidsinskii* (Vaneechoutte *et al*. 2019) Bouzek *et al*. comb. nov.

Basonym: *Gardnerella swidsinskii* Vaneechoutte *et al*. 2019^32^

*Gardnerella swidsinskii* (swid.sins’ki.i. N.L. gen. masc. n. *swidsinskii*, of Swidsinski, named after Alexander Swidsinski, a German microbiologist, who recognized that clue cells in Gram stains of vaginal swabs of women with BV are indicative for biofilm formation by *Gardnerella*).

The description of this subspecies remains as provided by Vaneechoutte *et al*. for *Gardnerella swidsinskii*^32^ with the following additional information. We included 46 genomes of cultivated *Gardnerella leopoldii* subsp. *swidsinskii* in our curated collection, isolated from the vagina, endometrium, rectum, and catheter urine of humans from China, France, Russia, and the United States (Supplementary Table 2). The median genome length is 1.63 Mb [range 1.52 – 1.72], and the median G+C content is 41.93% [range 41.64% - 42.29%]. The type strain of *Gardnerella leopoldii* subsp. *swidsinskii* remains GS 9838-1^T^ (=LMG 30812^T^=CCUG 72429^T^).

Description of *Gardnerella leopoldii* subsp. *aggregata* subsp. nov.

*Gardnerella leopoldii* subsp. *aggregata* (ag.gre.ga’ta. L. fem. part. adj. *aggregata*, joined together, referring to the presence of genes related to cell aggregation).

The genomes of *Gardnerella leopoldii* subsp. *aggregata*, as well as those of the overarching species, differ from other *Gardnerell*a species in the presence of genes consistent with adherence and biofilm formation. There are five genomes of cultivated *Gardnerella leopoldii* subsp. *aggregata* in our curated collection, isolated from the vagina and ectocervical mucosa of humans from Bangladesh, South Africa the United States, and Zambia (Supplementary Table 2). The median genome length is 1.48 Mb [range 1.48 – 1.53], and the median G+C content is 42.19% [range 42.11% - 42.50%]. This isolate is a gram variable coccobacillus and produces pinpoint colonies (≤ 1 mm) on Brucella agar after three days of anaerobic incubation at 37°C. Colonies are glossy white, round, convex, and alpha-hemolytic. In a genome comparison of *Gardnerella leopoldii* subsp. *aggregata* strain C0026F5^T^ with validly published *Gardnerella* type strains and the type strains proposed in this study, the highest ANI values correspond with its sister subspecies *Gardnerella leopoldii* subsp. *phocaeensis* Marseille-Q9179^T^ (96.09%), *Gardnerella leopoldii* subsp. *leopoldii* UGent 06.41^T^ (95.58%), and *Gardnerella leopoldii* subsp. *swidsinskii* GS 9838-1^T^ (94.70%), followed by *Gardnerella greenwoodii* subsp. *bretellae* Marseille-QA0894^T^ (87.40%) (Figure 2). Within the genome, 74 CAZymes were identified, including α-glucosidase (GH65), endo-β-galactosidase (GH16), and β-glucosidase (GH3)^112^. KEGG analysis identified a total of 84 transporters, 14 secretion genes, and 353 enzymes^112^. Metabolism predictions using GapMind^103–105^ include biosynthesis of glutamine, glycine, methionine, proline, and threonine, and catabolism of ethanol (Supplementary Table 6). The type strain of *Gardnerella leopoldii* subsp. *aggregata* is C0026F5^T^, isolated from the vagina of an asymptomatic menstruating woman from Birmingham, Alabama, USA. Accession for the 16S rRNA gene sequence for C0026F5^T^ is PZ016853.

Description of *Gardnerella leopoldii* subsp. *phocaeensis* (Allini Ntiguemassa *et al*. 2026) Bouzek *et al*. comb. nov.

Basonym: *Gardnerella phocaeensis* Allini Ntiguemassa *et al*. 2026^49^

*Gardnerella phocaeensis* (pho.cae.en’sis. L. fem. adj. *phocaeensis*, referring to Phocaea, the name of the Ionian Greek city where the founders of Marseille came from. The strain was isolated in Marseille).

The description of *Gardnerella leopoldii* subsp. *phocaeensis* remains as given to *Gardnerella phocaeensis* by Allini Ntiguemassa *et al*.^49^ with the following additional information. There are eleven genomes of cultivated *Gardnerella leopoldii* subsp. *phocaeensis* in our curated collection, isolated from the vagina and endometrium of humans from Bangladesh, France, the United States, and Zambia (Supplementary Table 2). The median genome length of *Gardnerella leopoldii* subsp. *phocaeensis* is 1.54 Mb [range 1.52 – 1.66] and the median G+C content is 42.26% [range 42.03% - 42.43%]. The type strain of *Gardnerella leopoldii* subsp. *phocaeensis* remains Marseille-Q9179^T^ (=CSUR Q9179^T^=CECT 31120^T^).

Description of *Gardnerella greenwoodii* subsp. *greenwoodii* (Sousa *et al*. 2023) Bouzek *et al.* comb. nov.

Basonym: *Gardnerella greenwoodii* Sousa *et al.* 2023^48^

*Gardnerella greenwoodii* (green.woo’di.i. N.L. gen. masc. n. *greenwoodii*, of Greenwood, named after James Greenwood, an American microbiologist who proposed the transfer of *Haemophilus vaginalis* to the new genus *Gardnerella*).

The description of this subspecies remains as given to *Gardnerella greenwoodii* by Sousa *et al*.^48^ with the following additional information. We included 13 genomes of *Gardnerella greenwoodii* subsp. *greenwoodii* in our curated collection, cultivated from the vagina, endometrium, and urine of humans from France, Portugal, the United Kingdom, and the United States (Supplementary Table 2). The median genome length of *Gardnerella greenwoodii* subsp. *greenwoodii* is 1.54 Mb [range 1.49 – 1.65] and the median G+C content is 43.40% [range 42.90% - 43.44%]. The type strain of *Gardnerella greenwoodii* subsp. *greenwoodii* remains c31Ua_26^T^ (=DSM 113415^T^=CCP 72^T^).

Description of Gardnerella greenwoodii subsp. intermedia subsp. nov.

*Gardnerella greenwoodii* subsp. *intermedia* (in.ter.me’di.a. L. fem. adj. *intermedia*, intermediate, between, referencing the shared possession of operons with both *Gardnerella* subgenera).

Pangenome analysis revealed that *Gardnerella greenwoodii* subsp. *intermedia*, as well as other members of the species, shares biologically relevant operons with the Set A subgenus (*Gardnerella vaginalis*, *Gardnerella piotii* and *Gardnerella rara*) while also falling within the Set B subgenus, suggesting that the species is an intermediate between Set A and Set B. Our curated set of genomes contains two strains of *Gardnerella greenwoodii* subsp. *intermedia*, both cultivated from the vagina of humans from the United States (Supplementary Table 2). The lengths of these genomes are 1.53 and 1.55 Mb, and the G+C contents are 43.04% and 43.05%. Cells are gram variable. Optimal growth occurs within two to three days on Brucella agar in an anerobic atmosphere at 37°C but can occur in aerobic environments. In a genome comparison of *Gardnerella greenwoodii* subsp. *intermedia* strain KA00603^T^ with validly published *Gardnerella* type strains and the type strains proposed in this study, the highest ANI values correspond with sister subspecies *Gardnerella greenwoodii* subsp. *bretellae* Marseille-QA0894^T^ (95.76%) and *Gardnerella greenwoodii* subsp. *greenwoodii* c31Ua_26^T^ (95.04%), followed by *Gardnerella kenyensis* NR047^T^ (88.32%) (Figure 2).

Within the genome, 66 CAZymes were identified, including β-glucosidase (GH3), and KEGG analysis identified a total of 85 transporters, 13 secretion genes, and 347 enzymes^112^. Metabolism predictions using GapMind^103–105^ include biosynthesis of cysteine, glutamine, glycine, methionine, proline, and threonine, and catabolism of ethanol and serine (Supplementary Table 6). The type strain of *Gardnerella greenwoodii* subsp. *intermedia* is KA00603^T,^ isolated from the vaginal fluid of a woman positive for bacterial vaginosis in Seattle, WA, USA. Accessions for the 16S rRNA gene sequence and whole genome sequence for KA00603^T^ are PZ013893 and CP058371.

Strain used in previous classifying publications: 1500E.

Description of *Gardnerella greenwoodii* subsp. *bretellae* (Allini Ntiguemassa *et al*. 2026) Bouzek *et al*. comb. nov.

Basonym: *Gardnerella bretellae* Allini Ntiguemassa *et al*. 2026

*Gardnerella bretellae* (bre.tel’lae. N.L. gen. fem. n. *bretellae*, honoring Florence Bretelle for her contribution to the description of vaginal microbiota).

The description of *Gardnerella greenwoodii* subsp. *bretellae* remains as given to *Gardnerella bretellae* by Allini Ntiguemassa *et al*.^49^ with the following additional information. Our curated collection of genomes contains eleven strains of *Gardnerella greenwoodii* subsp. *bretellae*, cultivated from the vagina and ectocervical mucosa of humans from Canada, France, Kenya, South Africa, the United States, and Zambia (Supplementary Table 2). The median genome length is 1.52 Mb [range 1.50 – 1.64], and the median G+C content is 43.22% [range 43.05% - 43.50%]. The *Gardnerella greenwoodii* subsp. *bretellae* type strain remains Marseille-QA0894^T^ (=CSUR QA0894^T^=CECT 31122^T^).

Description of Gardnerella washingtoni sp. nov.

*Gardnerella washingtoni* (wa.shing.to’ni. N.L. gen. n. *washingtoni*, named in honor of Washington University in St. Louis, MO, USA, where the first two genomes of this species originated, and the University of Washington in Seattle, WA, USA, which is associated with the first complete genome of this species).

There are five genomes of *Gardnerella washingtoni* in our curated collection. These isolates have been cultivated from the vagina and urine of humans in the United States (Supplementary Table 2). The median genome length of *Gardnerella washingtoni* is 1.60 Mb [range 1.56 – 1.65] and the median G+C content is 42.96% [range 42.89% – 43.11%]. Cells are Gram variable. The isolate is a facultative anaerobe that grows on Brucella agar in anaerobic conditions at 37°C in one to two days. In a genome comparison of *Gardnerella washingtoni* strain DNF00536^T^ with validly published *Gardnerella* type strains and the type strains proposed in this study, *Gardnerella greenwoodii* subsp. *bretellae* Marseille-QA0894^T^ has the highest ANI value (87.40%) (Figure 2).

Within the genome, 69 CAZymes were identified, including endo-β-galactosidase (GH16), α-glucosidase (GH65), and β-glucosidase (GH3)^112^. KEGG analysis identified a total of 83 transporters, 13 secretion genes, and 348 enzymes^112^. Metabolism predictions using GapMind^103–105^ include biosynthesis of cysteine, glutamine, glycine, methionine, and threonine, and catabolism of ethanol (Supplementary Table 6). The type strain of *Gardnerella washingtoni* is DNF00536^T^, isolated from the vaginal fluid of a woman positive for bacterial vaginosis in Seattle, WA, USA. Accessions for the 16S rRNA gene sequence and whole genome sequence for DNF00536^T^ are PZ013897 and CP058398.

Strains used in previous classifying publications: JCP8481A, JCP8481B, and PSS7772B.

Description of *Gardnerella bivia* sp. nov.

*Gardnerella bivia* (bi’vi.a. L. fem. adj. *bivia*, having two ways, in reference to the genome containing genes for two major pathways for mucin degradation).

*Gardnerella bivia* has genes associated with two major pathways for mucin degradation. There are six isolates of this species in our curated collection, cultivated from the vagina of humans in Bangladesh, the United States, and Zambia (Supplementary Table 2). The median genome length of *Gardnerella bivia* is 1.59 Mb [range 1.55 – 1.64] and the median G+C content is 38.08% [range 37.90% – 38.31%]. The isolate is a gram variable obligate anaerobe. Growth occurs on Brucella agar within one to two days in anaerobic conditions at 37°C. In a genome comparison of *Gardnerella bivia* strain DNF00502^T^ with validly published *Gardnerella* type strains and the type strains proposed in this study, *Gardnerella hutchinsoni* KA00225^T^ has the highest ANI value (82.91%) (Figure 2). Within the genome, 75 CAZymes were identified, including β-galactoside phosphorylase (GH112), α-L-fucosidase (GH29), and β-glucosidase (GH3)^112^. KEGG analysis identified a total of 74 transporters, 13 secretion genes, and 349 enzymes^112^. Metabolism predictions using GapMind^103–105^ include biosynthesis of cysteine, glutamine, glycine, proline, and threonine, and catabolism of acetate, ethanol, and serine (Supplementary Table 6). The type strain of *Gardnerella bivia* is DNF00502^T^, isolated from the vaginal fluid of a woman positive for bacterial vaginosis in Seattle, WA, USA. Accessions for the 16S rRNA gene sequence and whole genome sequence for DNF00502^T^ are PZ013890 and CP058399.

Strain used in previous classifying publications: CMW7778B.

Description of Gardnerella hutchinsoni sp. nov.

*Gardnerella hutchinsoni* (hutch.in.so’ni. N.L. gen. n. *hutchinsoni*, named in honor of Fred Hutchinson Cancer Center, where the isolate was first cultivated and the first complete genomes of this subspecies were sequenced).

We included six genomes of *Gardnerella hutchinsoni* isolates in our curated collection, all cultivated from the vagina of humans in the United States and Zambia (Supplementary Table 2). The median genome length of *Gardnerella hutchinsoni* is 1.67 Mb [range 1.64 – 1.72] and the median G+C content is 40.64% [range 40.53% – 40.76%]. Cells are gram variable. Optimal growth occurs within one to two days on Brucella agar in an anerobic atmosphere at 37°C but can occur in aerobic environments. In a genome comparison of *Gardnerella hutchinsoni* strain KA00225^T^ with validly published *Gardnerella* type strains and the type strains proposed in this study, *Gardnerella leopoldii* subsp. *aggregata* C0026F5^T^ has the highest ANI value (86.69%) (Figure 2). Within the genome, 69 CAZymes were identified, including β-galactoside phosphorylase (GH112) and β-glucosidase (GH3)^112^. KEGG analysis identified a total of 70 transporters, 13 secretion genes, and 341 enzymes^112^. Metabolism predictions using GapMind^103–105^ include biosynthesis of asparagine, glutamine, glycine, methionine, and proline, and catabolism of ethanol (Supplementary Table 6). The type strain of *Gardnerella hutchinsoni* is KA00225^T^, isolated from the vaginal fluid of a woman positive for bacterial vaginosis in Seattle, WA, USA. Accessions for the 16S rRNA gene sequence and whole genome sequence for KA00225^T^ are PZ013891 and CP058376.

Strains used in previous classifying publications: 3-005-3V and HUZ601990-0.

Description of Candidatus *Gardnerella kenyensis* sp. nov.

*Gardnerella kenyensis* (ken.yen’sis. N.L. fem. adj. *kenyensis*, from Kenya, referring to the country of isolation of the type strain).

All known strains of Candidatus *Gardnerella kenyensis* to date (NR002, NR021, and NR047) have been isolated from the vaginal fluid of a human in Nairobi, Kenya (Supplementary Table 2). Two of these are genomic duplicates (NR002 and NR047) and one (NR021) did not pass our genome quality threshold metrics. As such, only NR047^T^ was included in our curated collection (Supplementary Table 2). The genome length of this isolate is 1.66 Mb, and its G+C content is 43.62%. In a genome comparison of Candidatus *Gardnerella kenyensis* strain NR047T with validly published *Gardnerella* type strains and the type strains proposed in this study, the highest ANI values correspond with *Gardnerella insolita* subsp. *libera* C0085C4^T^ (90.59%) (Figure 2). Within the genome, 66 CAZymes were identified, including sialidase (GH33), and KEGG analysis identified a total of 84 transporters, 13 secretion genes, and 340 enzyme^112^. Metabolism predictions using GapMind^103–105^ include biosynthesis of asparagine, cysteine, glutamine, glycine, methionine, proline, and threonine, and catabolism of ethanol (Supplementary Table 6). The type strain of Candidatus *Gardnerella kenyensis* is NR047^T^ and the accession for the whole genome sequence is NQOH00000000.2.

Description of *Gardnerella insolita* sp. nov.

*Gardnerella insolita* (in.so’li.ta. L. fem. adj. *insolita*, unusual, strange, uncommon, referring to the low number of species cultivated).

To date, only three strains of *Gardnerella insolita* have been cultivated, all isolated from the vagina of humans in the United States and Zambia (Supplementary Table 2). The median genome length of *Gardnerella insolita* is 1.61 Mb [range 1.59 – 1.66] and the median G+C content is 43.60% [range 41.27% – 43.80%]. The *Gardnerella insolita* type strain is C0396A3^T^, isolated from the vagina of a pregnant woman at 19 weeks of gestation and who delivered preterm in Zambia. Accession for the 16S rRNA gene sequence C0396A3^T^ is PZ236487.

Description of *Gardnerella insolita* subsp. *insolita* comb. nov.

Basonym: *Gardnerella insolita* Bouzek *et al*. 2026

*Gardnerella insolita* (in.so’li.ta. L. fem. adj. *insolita*, unusual, strange, uncommon, referring to the low number of species cultivated).

There are two strains of *Gardnerella insolita* subsp. *insolita* in our curated genome collection. The genome lengths of these isolates are 1.66 and 1.61 Mb, and the G+C contents are 41.27% and 43.80%. This isolate is a gram-positive bacillus and produces pinpoint (≤ 1 mm) colonies on Brucella agar after two days of anaerobic incubation at 37°C. Colonies are round, clear, and glossy. In a genome comparison of *Gardnerella insolita* subsp. *insolita* strain C0396A3^T^ with validly published *Gardnerella* type strains and the type strains proposed in this study, the highest ANI values correspond with *Gardnerella insolita* subsp. *libera* C0085C4^T^ (95.24%), followed by *Gardnerella kenyensis* NR047^T^ (89.89%) (Figure 2). Within the genome, 66 CAZymes were identified, including β-glucosidase (GH3)^112^. KEGG analysis identified a total of 85 transporters, 13 secretion genes, and 347 enzymes^112^. Metabolism predictions using GapMind^103–105^ include biosynthesis of asparagine, glutamine, glycine, proline, and threonine, and catabolism of ethanol (Supplementary Table 6).

Strain used in previous classifying publications: 1-004-1.

Description of *Gardnerella insolita* subsp. *libera* subsp. nov.

*Gardnerella insolita* subsp. *libera* (li’be.ra. L. fem. adj. *libera*, independent, free, referring to the rarity of the subspecies).

To date, only one strain of *Gardnerella insolita* subsp. *libera* has been cultivated, C0085C4^T^, from the vagina of a woman positive for *Chlamydia trachomatis* in Baltimore, Maryland, USA. Its genome length is 1.59 Mb, and its G+C content is 43.60%. This isolate is a gram variable coccobacillus and produces pinpoint colonies (≤1 mm) on Brucella agar after two days of anaerobic growth at 37°C. Colonies are grey, glistening, round, and β-hemolytic. In a genome comparison of *Gardnerella insolita* subsp. *libera* strain C0085C4^T^ with validly published Gardnerella type strains and the type strains proposed in this study, the highest ANI values correspond with *Gardnerella insolita* subsp. *insolita* C0396A3^T^ (95.30%), followed by Candidatus *Gardnerella kenyensis* NR047^T^ (90.82%) (Figure 2). Within the genome, 70 CAZymes were identified, including Class II α-mannosidase and α-glucosidase^112^. KEGG analysis identified a total of 84 transporters, 12 secretion genes, and 342 enzymes^112^. Metabolism predictions using GapMind^103–105^ include biosynthesis of cysteine, glutamine, glycine, proline, and methionine, and catabolism of ethanol and serine (Supplementary Table 6). The type strain of *Gardnerella insolita* subsp. *libera* is C0085C4^T^. Accession for the 16S rRNA gene sequence for C0085C4^T^ is PZ016852.

## Acknowledgements

Research reported in this publication was supported in part by the National Human Genome Research Institute as part of the Human Microbiome Project Initiative focused on technology development under award number R01HG005816 (D.N.F.), the National Institute for Allergy and Infectious Diseases of the National Institutes of Health under award number U19AI084044 (J.R.), the National Institute of Dental and Craniofacial Research (NIDCR) of the National Institutes of Health Office of the Director (OD) Common Fund under award number R01DE027850 (C.D.J). and the Gates Foundation under award INV010470 to (J.R.). In addition, startup funds were provided by Fred Hutch Cancer Center and MD Anderson Cancer Center (C.D.J.).

## Contributions

H.K.B., S.S., C.D.J., and D.N.F. conceived the study. E.F.M., D.S.J., S.M.S., T.F.L., E.M.L., and M.F. conducted experiments including strain isolation, DNA extraction, whole-genome sequencing, cloning, and testing of the shuttle vector. H.K.B. and M.Z.R. conducted all bioinformatics and curated data. S.S., C.D.J, J.R., and D.N.F. supervised and obtained funding for the project. H.K.B., M.Z.R., S.S., S.M.S., C.D.J, and D.N.F. wrote the original draft. J.R. and M.K. provided critical edits to the manuscript and assisted in nomenclature. M.K. provided crucial nomenclature feedback and provided edits to the protologues and manuscript. All authors were involved in reviewing and editing this submission.

## Data availability

NCBI Accession numbers of genomes used are available in Supplementary Table 1.

