## Supplementary Figures 1-11 for "A Syntenic Pangenome of *Gardnerella* Reveals Novel Plasmids and Phage, Taxonomic Boundaries, and Species-Level Stratification of Metabolic and Virulence Potential"

### Extended Data Figures

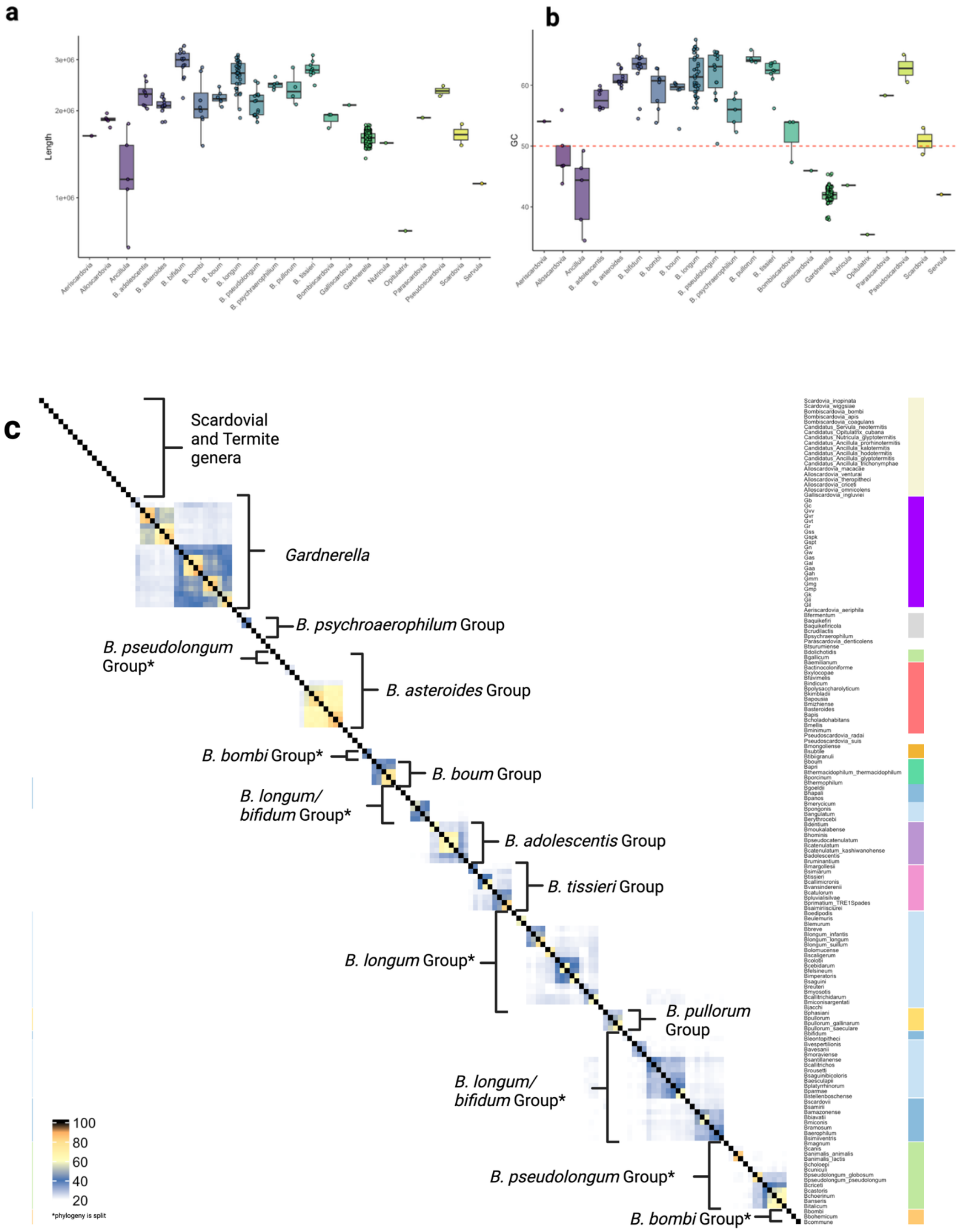

**d**

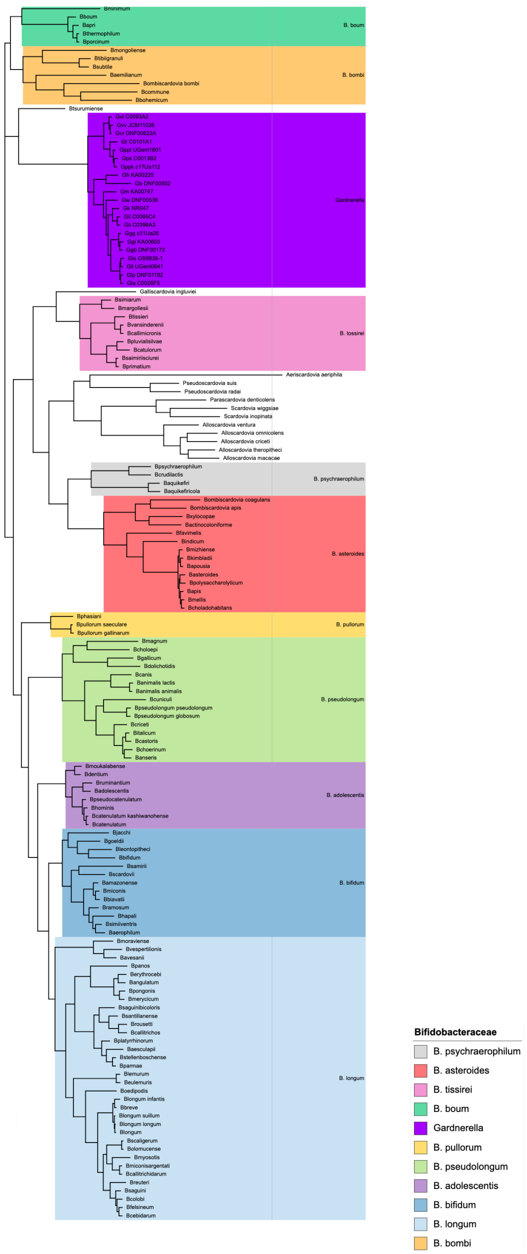
B. boum
B. bombi
Gardnerella
B. tissirei
B. psychraerophilum
B. asteroides
B. pullorum
B. pseudolongum
B. adolescentis
B. bifidum
B. longum
Bifidobacteraceae
B. psychraerophilum
B. asteroides
B. tissirei
B. boum
Gardnerella
B. pullorum
B. pseudolongum
B. adolescentis
B. bifidum
B. longum
B. bombi

e

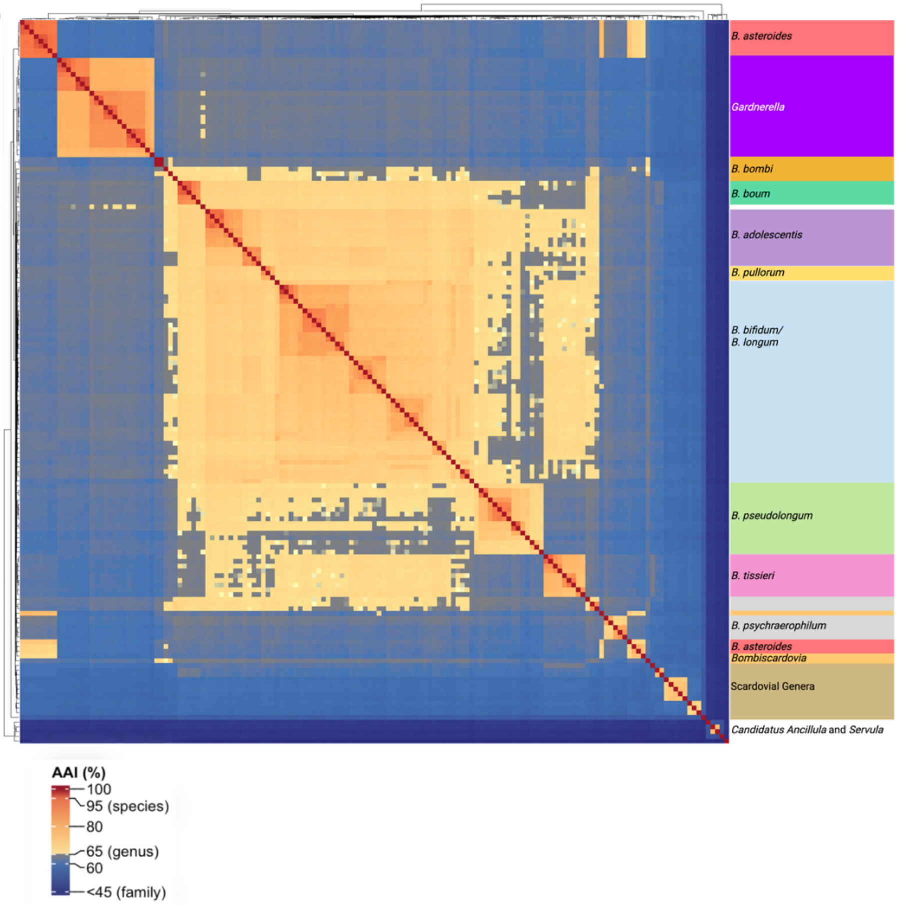

B. asteroides
Gardnerella
B. bombi
B. boum
B. adolescentis
B. pullorum
B. bifidum/
B. longum
B. pseudolongum
B. tissieri
B. psychraerophilum
B. asteroides
Bombiscardovia
Scardovial Genera
Candidatus Ancillula and Servula
AAI (%)
100
95 (species)
80
65 (genus)
60
<45 (family)

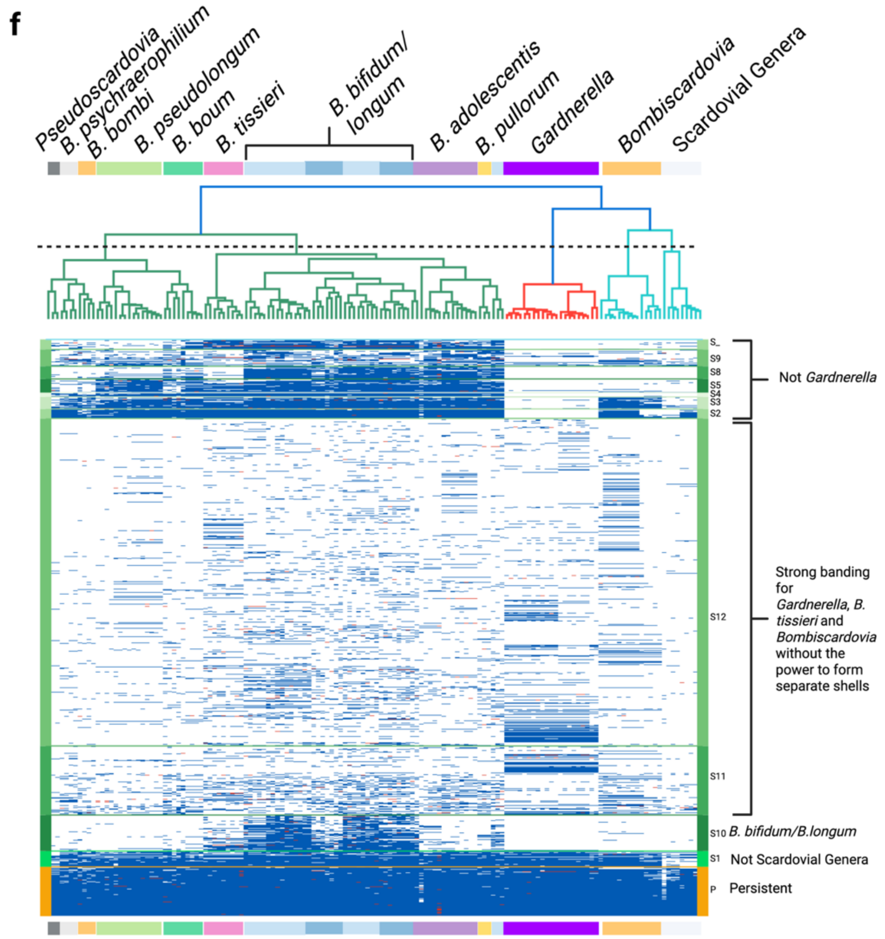


**Extended Data Fig. 1: Genomic comparisons among members of the family *Bifidobacteriaceae***

Comparative genomic analyses across representative genera of the family *Bifidobacteriaceae*, including subgroups of *Bifidobacterium* (as defined by Alessandri *et al.*), the "Scardovial" genera (*Aeriscardovia*, *Alloscardovia*, *Bombiscardovia*, *Galloscardovia*, etc.), and genera associated with termite gut microbiota (*Ancillula*, *Servulla*, etc.).

**a,** Genome length comparisons across representative genera. Relative to *Bifidobacterium*, genomes of *Gardnerella* (Kruskal–Wallis with Bonferroni correction, $p < 0.001$) and *Alloscardovia* ($p = 0.052$) are smaller. *Gardnerella* genomes are also significantly smaller than those of *Alloscardovia* ($p = 0.009$). Other comparisons involving Scardovial genera lack sufficient representation or exhibit broad variance that limits statistical inference. **b,** Genomic G+C content across selected genera. *Gardnerella*, *Ancillula*, and *Alloscardovia* exhibit significantly lower G+C percentages than *Bifidobacterium* ($p < 0.001$, $p = 0.0113$, and $p = 0.0179$, respectively).

**c,** Digital DNA–DNA hybridization (dDDH) values among representative genomes. The species-level threshold (>70%) is highlighted in yellow; intermediate similarity (~40%) is shown in blue, and low similarity (<20%) in white. *Gardnerella* forms a monophyletic cluster within *Bifidobacteriaceae*. Groups marked with an asterisk indicate polyphyletic lineages. Colors along the axes correspond to the *Bifidobacterium* groups defined by Alessandri *et al.*

**d**, Conserved protein phylogeny of *Bifidobacteriaceae* type strains. Persistent PPanGGOLiN gene clusters were extracted and used to construct a neighbor-joining tree. *Gardnerella* forms a monophyletic lineage within the broader *Bifidobacteriaceae*, with the nearest cluster including a member of *Bombiscardovia*.

**e,** Amino acid identity (AAI) among type strains. Pairwise AAI values were calculated and visualized as a heatmap. Color gradients correspond to commonly applied genomic thresholds: dark blue (<45%, typical of inter-family divergence), light blue (45–65%, inter-generic range), yellow (>65%, genus-level coherence), and red (~100%, near-identical comparisons).

**f,** PPanGGOLiN tile map of the pangenome derived from *Bifidobacteriaceae* type strains. Nodes (rows) correspond to gene clusters within the persistent fraction of the pangenome (orange bar). Shell partitions are indicated by green shades. The accompanying dendrogram is derived from a similarity matrix of gene presence–absence patterns. *Gardnerella* clusters with the Scardovial genera. The tile plot shows the relatively few persistent genes are uniquely shared between *Bifidobacterium* and *Gardnerella*.

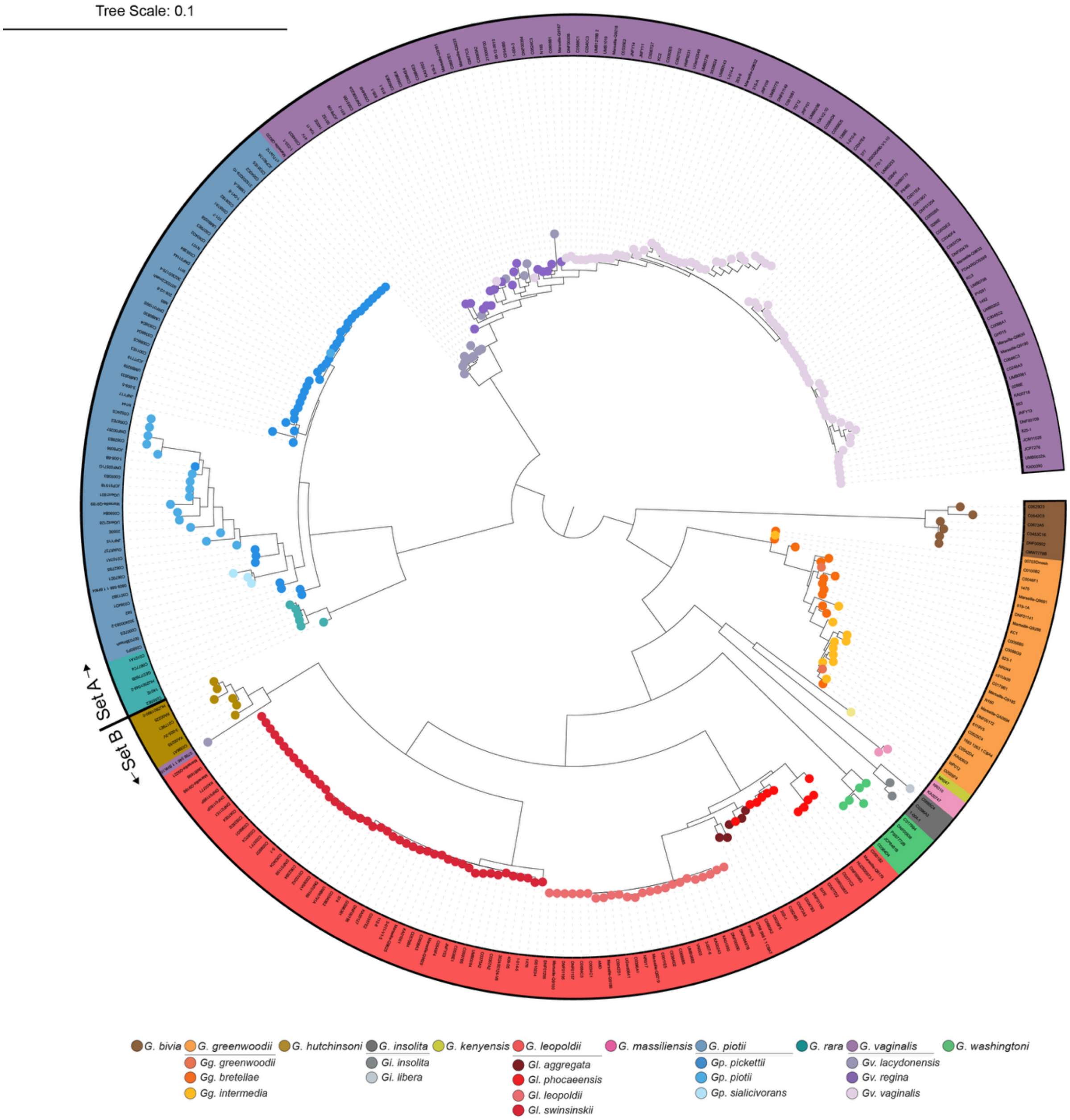


**Extended Data Figure 2: *Gardnerella cpn60* gene sequence dendrogram**

Maximum-likelihood dendrogram based on MUSCLE-aligned *cpn60* gene sequences from *Gardnerella* genomes. Label indicates strain identifier, while label and tip point color represent lineage.

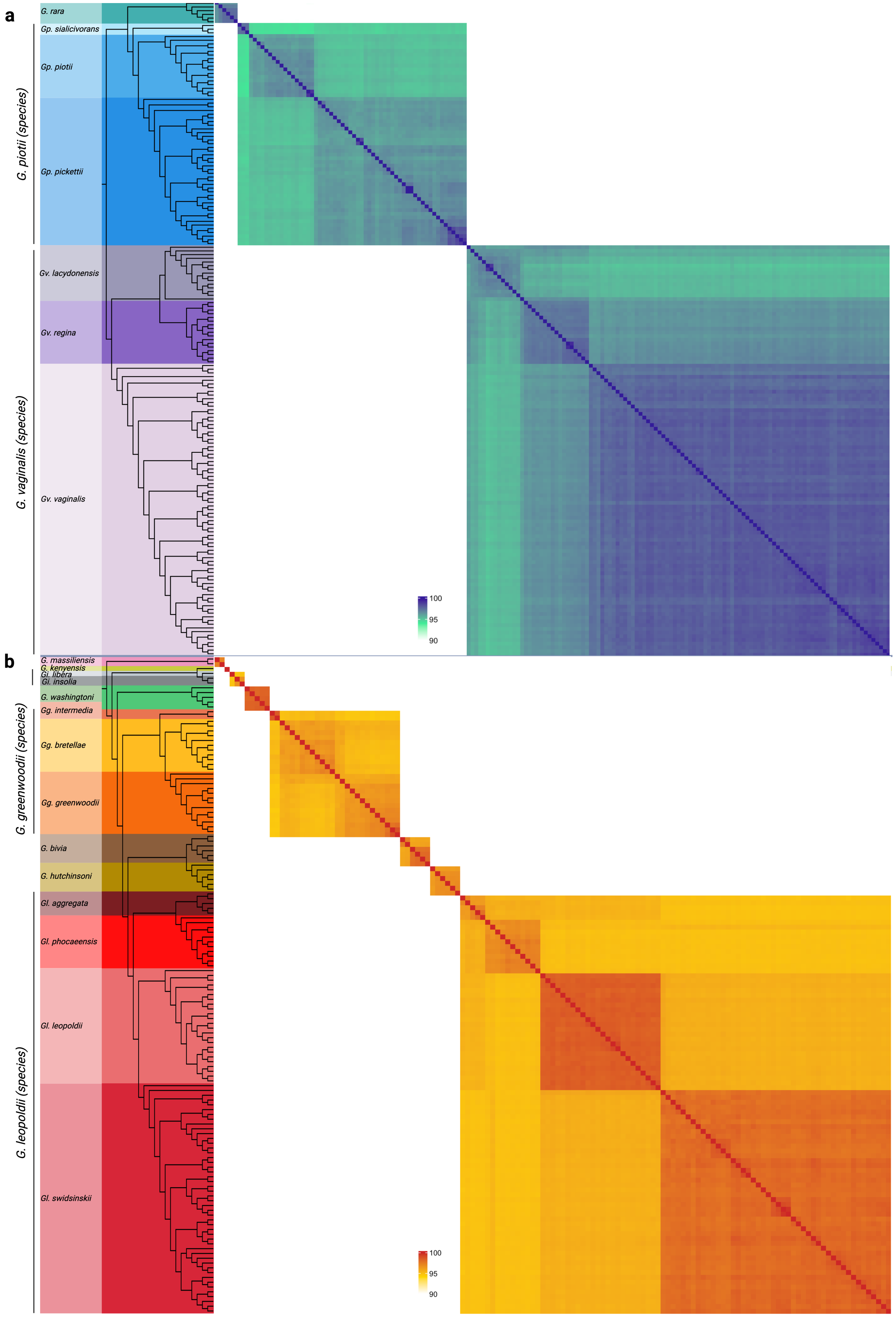
a
b
G. rara
Gp. sialicivorans
Gp. piotii
Gp. pickettii
G. piotii (species)
Gv. lacydonensis
Gv. regina
Gv. vaginalis
G. vaginalis (species)
100
95
90
G. massiliensis
G. kenyensis
Gi. libera
Gi. insolia
G. washingtoni
Gg. intermedia
Gg. bretellae
Gg. greenwoodii
G. greenwoodii (species)
G. bivia
G. hutchinsoni
Gl. aggregata
Gl. phocaeensis
Gl. leopoldii
Gl. swidsinskii
G. leopoldii (species)
100
95
90

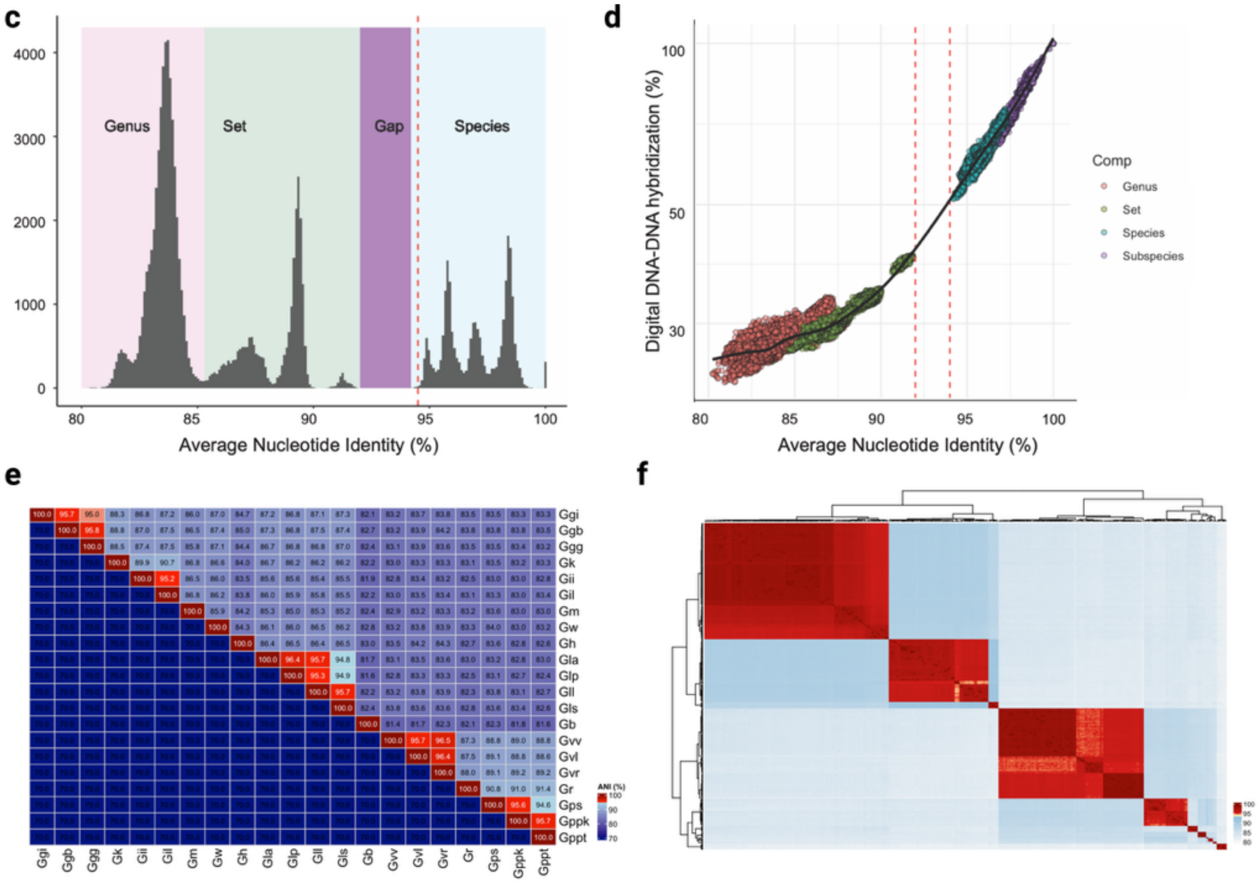
c
Genus
Set
Gap
Species
4000
3000
2000
1000
0
80
85
90
95
100
Average Nucleotide Identity (%)
d
Digital DNA-DNA hybridization (%)
100
50
30
Average Nucleotide Identity (%)
Comp
Genus
Set
Species
Subspecies
e
Ggi
Ggb
Ggg
Gk
Gii
Gil
Gm
Gw
Gh
Gla
Glp
Gll
Gls
Gb
Gvv
Gvl
Gvr
Gr
Gps
Gppk
Gppt
ANI (%)
100
90
80
70
f
100
95
90
85
80

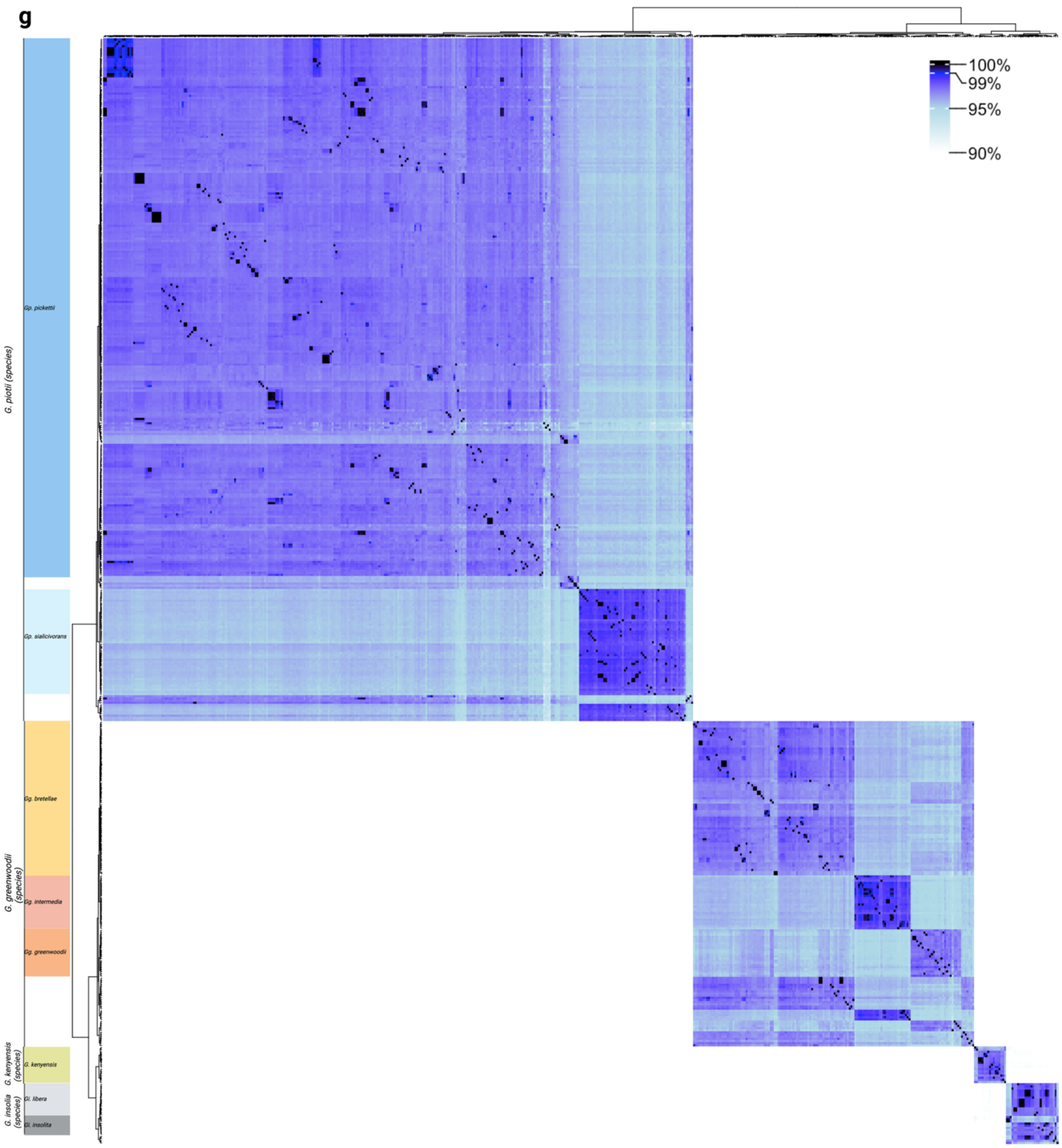


**Extended Data Figure 3: Average nucleotide identity (ANI) and digital DNA-DNA hybridization** (dDDH) comparisons between *Gardnerella* genomes. **a,** dDDH comparison of the complete collection of Set A genomes as ordered by kSNP phylogeny **b**, dDDH comparison of the complete collection of Set B genomes as ordered by kSNP phylogeny. **c,** A histogram of the counts of ANI percentage comparisons between *Gardnerella* genomes showing the genus, set (subgenus), and species comparisons at each level. The gap between species and higher order comparisons falls slightly lower than 95% (red dashed line). **d**, Collinearity of ANI and dDDH values with red dashed lines showing the boundaries of the gap in both values. **e,** ANI of the type strains of *Gardnerella* species and subspecies. The mean of the pairwise ANI (i,j) and (j,i) represented as just the upper triangle. Values from 100% to 95% are colored in increasingly lighter shades of red, values below 95.0% are colored in increasingly darker shades of blue. **f,** ANI of the *Gardnerella* strains in the curated collection with an emphasis on the division at 95% which is colored white. This emphasizes that the few strains that do fall in the 95-94% range are well within otherwise sharply defined species groups. **g**, Heatmap of pairwise ANI values for species in *Gardnerella* with low numbers of cultured isolates at the subspecies level as compared to MAGs (metagenome assembled genomes). Colored bands at the left give the species and subspecies groups for the boxes in the heatmap. Rows without coloration are to MAGs that are unassigned to species.

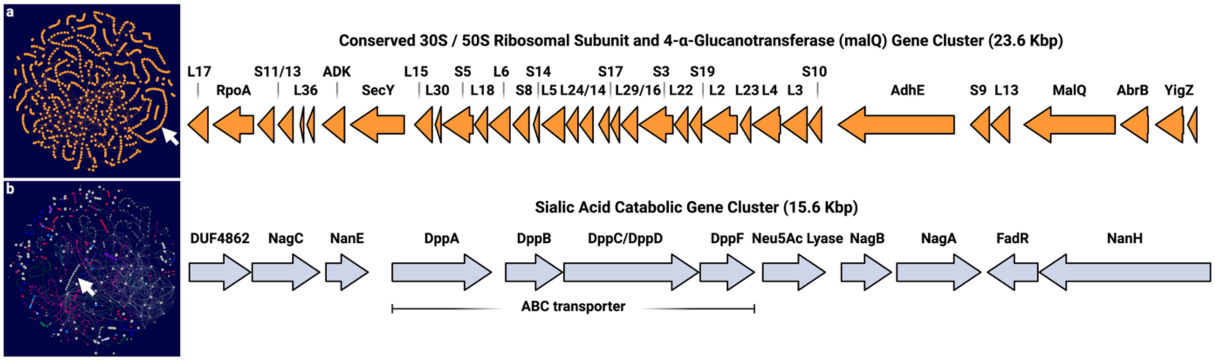


**Extended Data Fig. 4: PPanGGOLiN syntenic nodes create modules**

**a,** Core and b, shell partitions from PPanGGOLiN-generated *Gardnerella* pangenome. Each node represents a gene group where the size of the node indicates relative presence across *Gardnerella* genomes. White arrows pointing to syntenic nodes represent neighboring genes in operons, shown on the right.

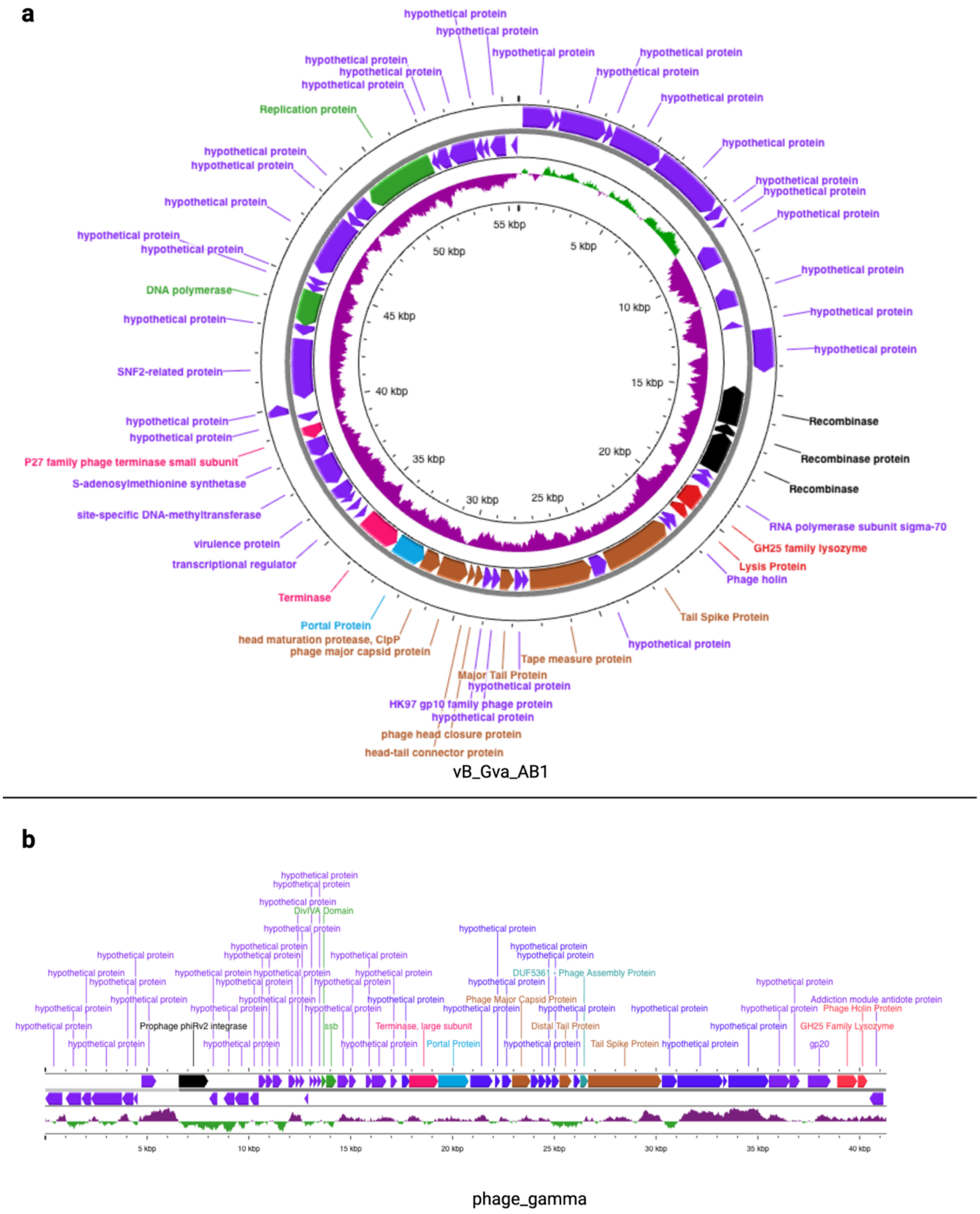
a
hypothetical protein
Replication protein
DNA polymerase
SNF2-related protein
P27 family phage terminase small subunit
S-adenosylmethionine synthetase
site-specific DNA-methyltransferase
virulence protein
transcriptional regulator
Terminase
Portal Protein
head maturation protease, ClpP
phage major capsid protein
head-tail connector protein
phage head closure protein
HK97 gp10 family phage protein
Major Tail Protein
Tape measure protein
Tail Spike Protein
Phage holin
Lysis Protein
GH25 family lysozyme
RNA polymerase subunit sigma-70
Recombinase
Recombinase protein
55 kbp
5 kbp
10 kbp
15 kbp
20 kbp
25 kbp
30 kbp
35 kbp
40 kbp
45 kbp
50 kbp
vB_Gva_AB1
b
hypothetical protein
DivIVA Domain
Prophage phiRv2 integrase
ssb
Terminase, large subunit
Portal Protein
DUF5361 - Phage Assembly Protein
Phage Major Capsid Protein
Distal Tail Protein
Tail Spike Protein
Addiction module antidote protein
Phage Holin Protein
GH25 Family Lysozyme
gp20
5 kbp
10 kbp
15 kbp
20 kbp
25 kbp
30 kbp
35 kbp
40 kbp
phage_gamma

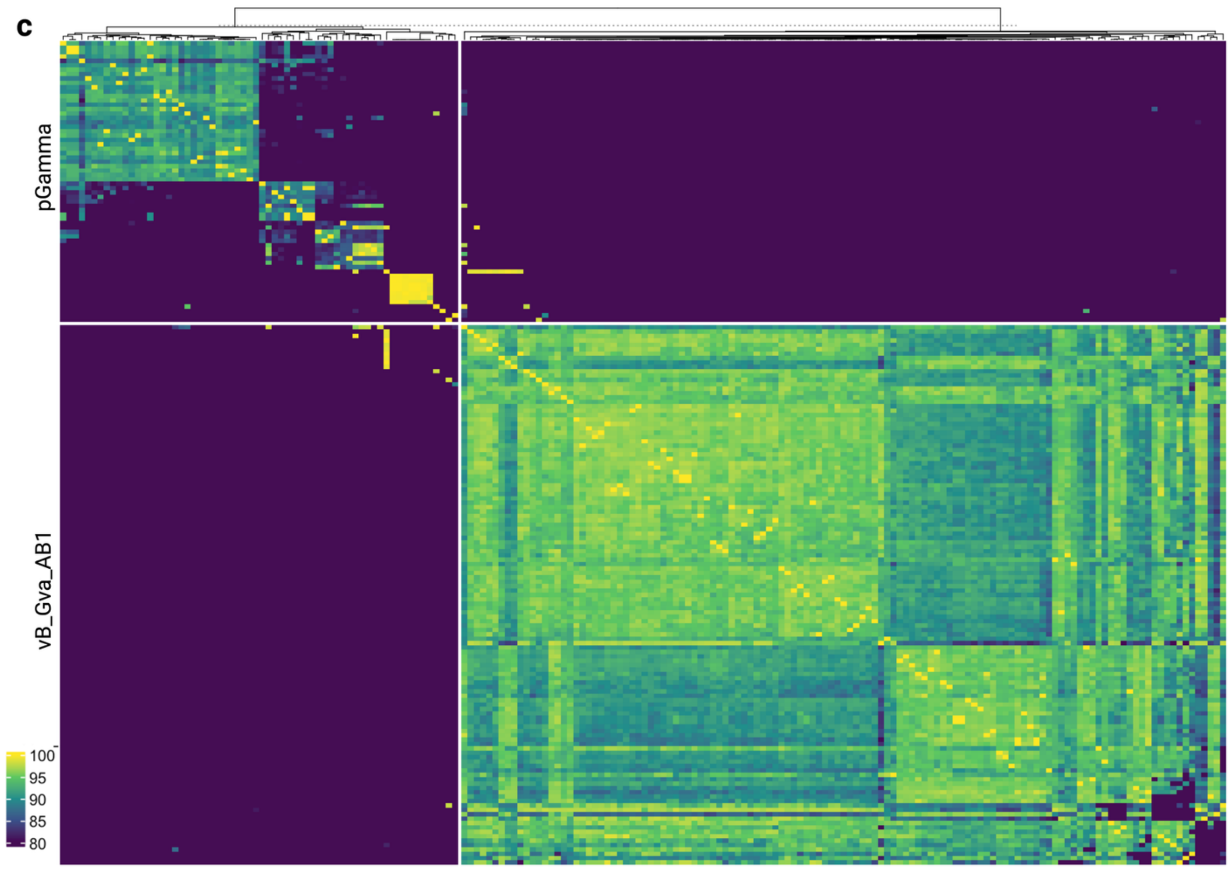


**Extended Data Figure 5: Bacteriophages in *Gardnerella***

Phage maps of the different families of bacteriophage found in *Gardnerella*. **a,** Example of the vB_Gva_AB1 phage found in circular lytic form in *Gardnerella* strain 06-12-0010 and the lysogenic form of pGamma found in DNF00622A in **b,** Recombination genes are black, structural genes are brown, lysis/terminase genes are bright red, portal genes are blue, assembly genes are in green. and hypothetical genes are purple. The inner ring/bottom of the graph shows the GC% with high GC as purple and above the line while green means low GC and is below the line. **c,** ANI of the complete phages found in this collection. Clustering was by km=2 for the two phage types. The largest block closely aligns to vB_Gva_AB1 phage. The smaller group made up of >3 smaller subgroups. The group that forms the strongest cohesion (yellow box in pGamma) are all from a single lab in China (Zhang 2022) and may overrepresent the natural occurrence of phage of this type.

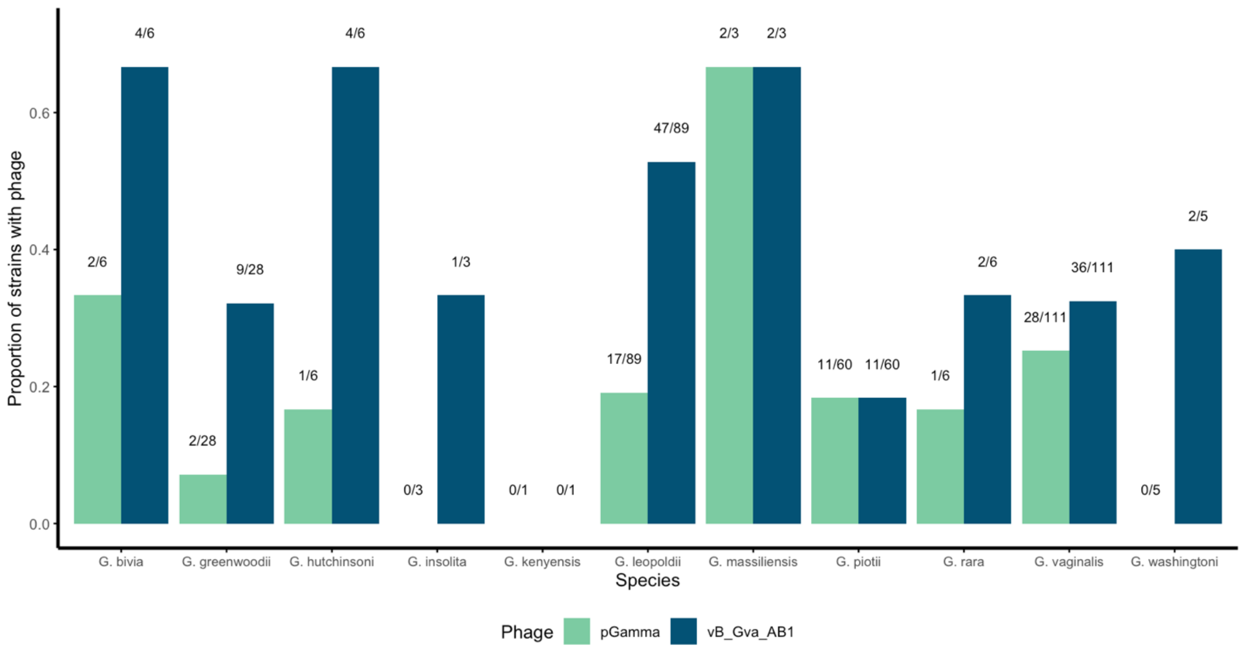


**Extended Data Figure 6: Phage Genomes**

Graphs show the proportion number of intact prophages detected in each species. Prophages were found alone or in pairs. The difference in proportion infected by phages of different types was significant for *G. leopoldii* (Bonferroni adj. $p < 0.01$, Fisher's Exact test) and *G. greenwoodii* (Bonferroni adj. $p = 0.4$, Fisher's Exact test). The differences in phage infection between sets and species was not significant (Chi Squared and Fisher's Exact tests, respectively).

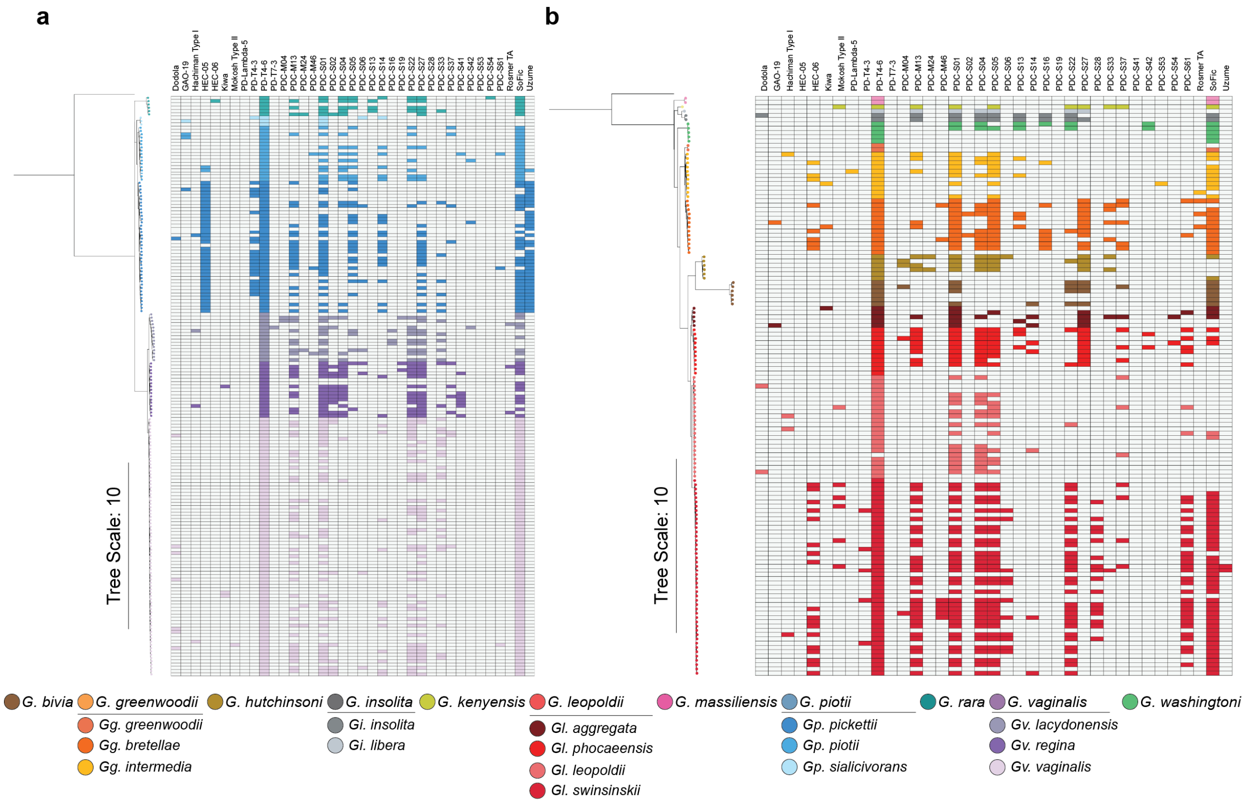


**Extended Data Figure 7: Predicted genomic defenses with undescribed mechanisms**
PADLOC-predicted defenses with currently undescribed mechanisms of action across *Gardnerella* genomes in **a,** Set A or **b,** Set B. Genomes are ordered by kSNP4 phylogenies, with colors indicating lineage.

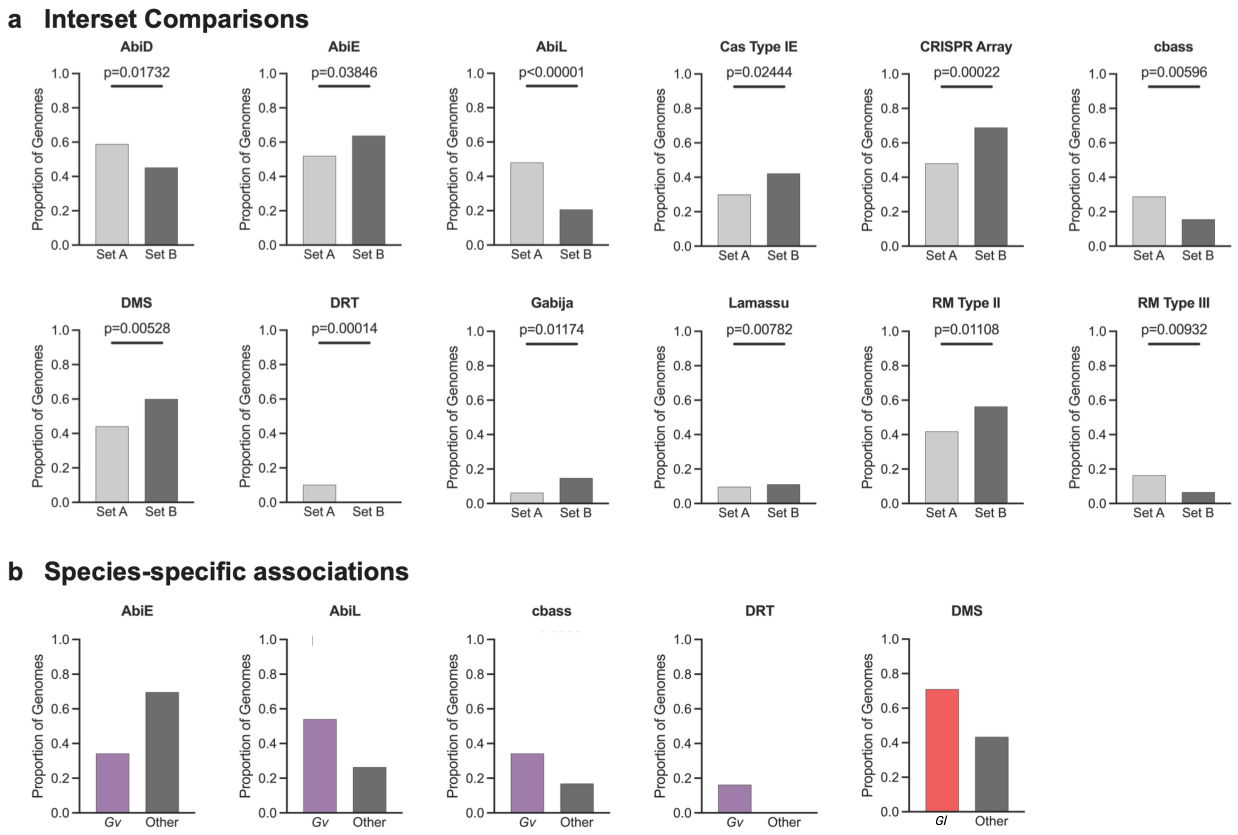


**Extended Data Figure 8: Genetic defense system associations across *Gardnerella* groups**

Graphs show **a,** the presence of PADLOC-predicted defenses compared by genome proportion between *Gardnerella* Set A and Set B and **b,** specific species, and all other species. Statistical analysis was carried out using a two-sample z-test, two-tailed.

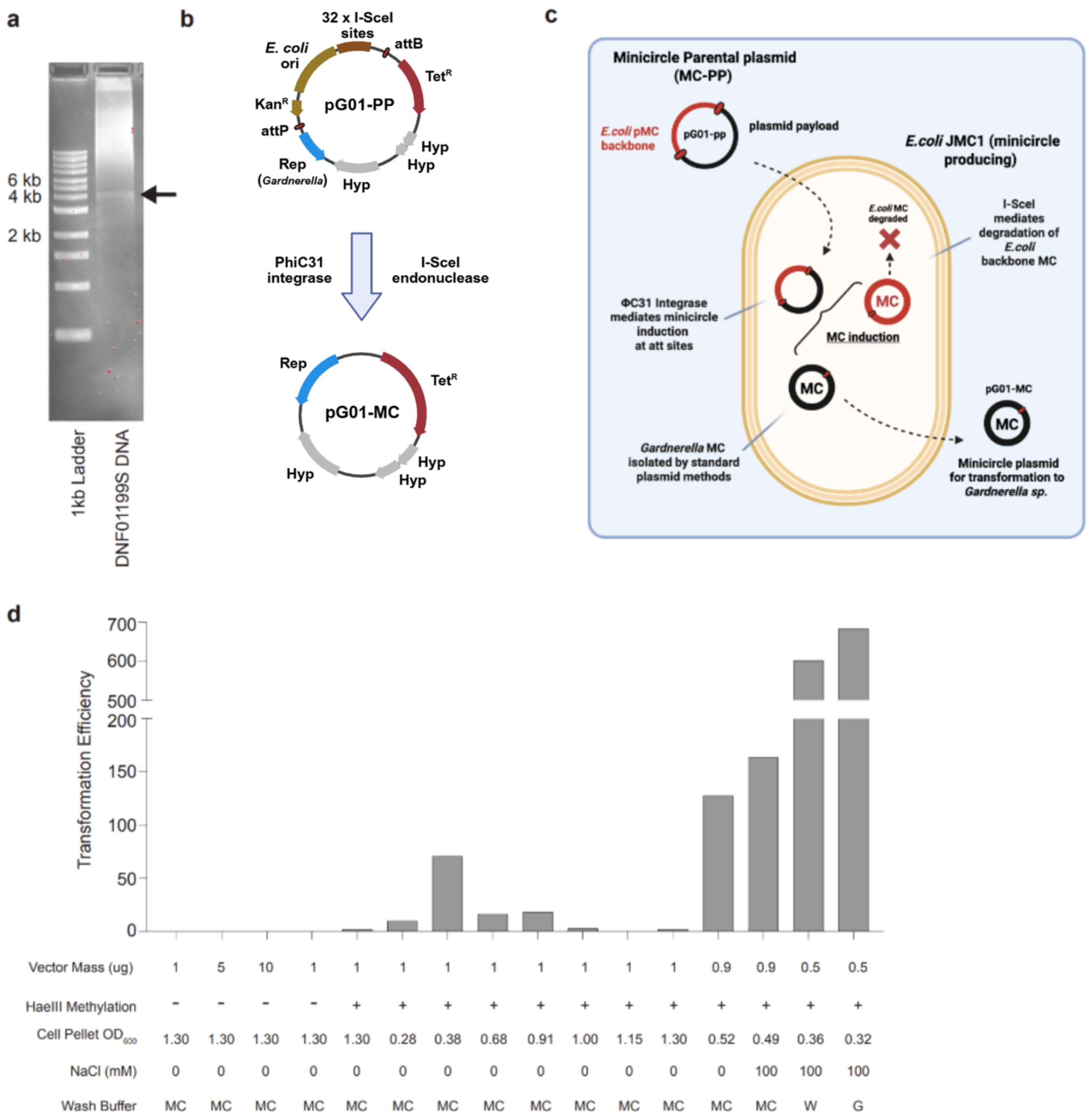


**Extended Data Figure 9: Transformation in *Gardnerella* strain ATCC 14018 of a vector tool derived from a native *Gardnerella* plasmid a,** pDNF0199S-01, a 4,028 bp plasmid, isolated from *Gardnerella* strain DNF01199S. **b-c,** Schematic shows the components of pG01-pp, **b,** the minicircle parental plasmid derived from pDNF01199S-01, and **c,** the resulting pG01-MC produced after minicircle induction. **d,** Transformation efficiency of pG01-MC in *Gardnerella* strain ATCC 14018 under different conditions, including changes to vector mass, the pre-methylation of pG01-MC with *HaeIII* to methyl-modify GGCC sites, the $OD_{600}$ of the cell pellet used, the concentration of sodium chloride (NaCl) present in the cell growth media, and the wash buffer used during electrocompetent cell preparation (MC=maltose citrate, W=milliQ water, G=10% glycerol).

#### Supplementary Table Legends

**Supplementary Table 1: Uncurated *Gardnerella* Genomes**

Table of the 649 *Gardnerella* genomes obtained via sequencing of Fredricks and Ravel isolate collections as well as the publicly-available genomes from the NCBI database. For each genome, available metadata includes BioSample, BioProject, Accession Name, Level, and Source, WGS Project Accession, Biological Source, Origin, Country, and Sequencing Method. For each genome, quality metrics include the number of contigs, genome length, GC content, and CheckM statistics. Genomic duplicates are noted.

**Supplementary Table 2: Curated *Gardnerella* Genomes**

Table of the 313 *Gardnerella* genomes as a final curated genome set. This set was generated by the removal of genomes with a genomic duplicate (an average nucleotide identity greater than 99.5%), CheckM contamination (greater than 1%), highly fragmented genomes (greater than 100 contigs), or completeness (less than 90%). Type strains were all retained and exempted from these conditions. The metadata available for each genome includes , as well as the *cpn60* gene sequence and BLAST results, and taxonomic classification via homology to the Genome Taxonomy Database using GTDB-tk is listed. Final taxonomic classification with the nomenclature proposed in this work is shown for each genome.

**Supplementary Table 3: *Bifidobacteriaceae* Lineages**

Table shows the 279 genomes from *Bifidobacterium*, *Aeriscardovia*, *Alloscardovia*, *Ancillula*, *Bombiscardovia*, *Galloscardovia*, *Optulatrix*, *Nutricula*, *Pseudoscardovia*, and *Scardovia* used for the comparative analyses in Supplementary Figure 1. The expanded Alessandri *Bifidobacterium* groups (Adding Scardovial and Termite Gut groups from within *Bifidobacteraceae*) and associated hexadecimal (HEX) codes used in the figures are given.

**Supplementary Table 4: *Gardnerella* Lineages**

Table shows the species and subspecies nomenclature proposed for each *Gardnerella* lineage identified in this work, as well as the range of ANI and dDDH values within each group. Included are the unique abbreviations and hexadecimal (HEX) color codes used throughout the figures and tables for these taxa.

**Supplementary Table 5: *Gardnerella* PPanGGOLiN Pangenome Dataset**

Table shows nodes (gene groups) identified by PPanGGOLiN pangenome analysis of the curated *Gardnerella* genome set. For each node, its product annotation, gene name, partition (persistent, shell, cloud), median length, the number of genomes it was identified in, and its module number (when applicable) are shown. Columns 7-318 show for each genome, the PGAP annotation identifier for the gene grouped into each node.

**Supplementary Table 6: GapMind Metabolic Predictions across *Gardnerella* Genomes**

Table shows the presence or absence of GapMind-predicted metabolic pathways for the biosynthesis of amino acids and the catabolism of small carbon sources across our curated *Gardnerella* genome set.

**Supplementary Table 7: Prophage Genes Detected Across *Gardnerella* Genomes**

Table shows the number of prophage mapped to either vB_Gva_AB1 or pGamma for each genome in our curated *Gardnerella* genome set as predicted by either PPanGGOLiN synteny and partitioning, VIBRANT, or PhageR. ANI types are assigned based on ANI homology in Extended Figure 5.

**Supplementary Table 8: PADLOC Genetic Defense System Predictions across *Gardnerella* genomes**
Table shows the presence or absence of PADLOC-predicted genetic defense systems across our curated *Gardnerella* genome set.

**Supplementary Table 9: Methyl-modified Motifs of a Subset of *Gardnerella* Genomes**
REBASE analysis of methyl-modified motifs detected during PacBio sequencing of *Gardnerella* isolates. REBASE organism accession numbers are listed in Supplementary Table 2.

**Supplementary Table 10: Nucleotide Sequences of *Gardnerella* Native Plasmids**
For the three native *Gardnerella* plasmids identified in our pangenome analysis, table shows plasmid name, length, and nucleotide sequence.

**Supplementary Table 11: Annotated Open Reading Frames in *Gardnerella* Native Plasmids**
Table shows the Proksee-predicted open reading frames, their corresponding amino acid sequence, top BLASTp hit, and conserved domain search results for each of the three native *Gardnerella* plasmids.
